# Laminin and Fibronectin contribute to Guided Endothelial Self-Organization During Intersegmental Vessel Formation

**DOI:** 10.64898/2026.03.13.711615

**Authors:** Joaqúın Abugattas-Núñez Del Prado, Koen A. E. Keijzer, Erika Tsingos, Roeland M. H. Merks

## Abstract

Endothelial cells can self-organize into vascular networks, but how tissue-derived cues constrain this behavior during stereotyped angiogenic sprouting is unclear. In zebrafish intersegmental vessels (ISVs), laminin and fibronectin are enriched in the intersomitic space. We integrated live imaging, perturbation experiments, rescue assays, and mathematical modeling to test whether these matrix components contribute to constraining endothelial network-forming behavior during ISV formation. Combined depletion of laminin alpha chains or fibronectin paralogs slowed endothelial sprouting without abolishing overall ISV formation. In a hybrid mathematical model coupling endothelial self-organization to a deformable extracellular matrix, simulations predicted that reducing matrix stiffness or density would slow sprouting and increase lateral vessel fusion. Consistent with this qualitative prediction, combined morpholino knockdown of *lama1*/*lama4*/*fn1a* (MO L14Fa) or *lama1*/*lama4*/*fn1b* (MO L14Fb) produced severe ISV mispatterning, including ectopic branching, abnormal inter-ISV connections, and network-like vascular arrangements; direct vessel fusion was captured by time-lapse imaging. Chimeric fibronectin mRNAs rescued vascular patterning in the corresponding combined perturbation backgrounds. Because matrix perturbation also disrupted trunk and somite organization, we considered whether secondary disruption of semaphorin guidance could explain the vascular defects. However, hybridization chain reaction (HCR) showed that *sema3aa* and *sema3ab* remained expressed in the trunk, with a relatively preserved *sema3ab* pattern, while RT-qPCR detected no significant differences in their overall transcript abundance; only local differences in *sema3aa* accompanied the more severe tissue disruption in MO L14Fa. Together with the comparison of *lama1*/*lama4*/*fn1a* and *lama1*/*lama4*/*fn1b* morphants, these findings suggest that trunk and somite abnormalities can exacerbate ISV defects but do not fully account for the vascular phenotype. Overall, our results support a guided self-organization model in which laminin and fibronectin, together with other tissue-derived cues, channel an intrinsic endothelial capacity for network formation toward stereotyped intersegmental sprouts. More broadly, our findings suggest that vasculogenic network formation and angiogenic sprouting may draw on overlapping endothelial self-organizing mechanisms whose outcomes depend on initial tissue configuration and environmental constraints.

## 1 Introduction

Angiogenesis, the formation of new vessels by sprouting from existing vasculature, and vasculogenesis, the assembly of endothelial cells into vascular networks, are conventionally distinguished by the origin of the newly formed vessels. During vasculogenesis, initially dispersed endothelial cells migrate, connect, and self-organize into network-like structures, a capacity also observed under permissive conditions *in vitro* (*1, 2, 3*). Induced zebrafish endothelial cells retain this network-forming capacity (*4*), indicating that it is not restricted to endothelial cells from a particular developmental or tissue context. Angiogenic sprouts also display self-organizing collective behaviors: endothelial cells dynamically rearrange and exchange positions within sprouts, form branches, and connect through anastomosis (*5, 6, 7, 8, 9*). Thus, although angiogenesis and vasculogenesis differ in their initial tissue configuration and resulting geometry, both draw on a shared repertoire of dynamic endothelial interactions. This raises the possibility that, at the level of collective cell behavior, angiogenesis and vasculogenesis draw on overlapping self-organizing mechanisms despite their distinct developmental contexts. Mathematical models provide mechanistic support for exploring this possibility. A broad range of models reproduce endothelial network formation using a small number of experimentally supported local behaviors (*10, 11, 12, 13, 14, 15, 16, 17, 18*). Although their implementations differ, a recurring principle is an attractive interaction between endothelial cells, mediated by chemotaxis, direct cellular pulling, or cell-generated mechanical forces, combined with a mechanism that prevents their collapse into compact aggregates. Notably, models based on the same local interactions can generate both networks from dispersed endothelial cells and sprouts from pre-existing endothelial structures (*13, 19*). This does not imply that angiogenesis and vasculogenesis are the same process. Rather, it suggests a narrower mechanistic possibility: endothelial cells may use overlapping local interaction rules in both contexts, while different initial configurations and environmental constraints produce different geometries.

The formation of intersegmental vessels (ISVs) in zebrafish (*Danio rerio*) provides a tractable system in which to test this view. During trunk vascular development, ISVs sprout dorsally from the dorsal aorta toward VEGF signals from the ventral somites, elongate along the intersomitic boundaries, and subsequently anastomose to form the dorsal longitudinal anastomotic vessel (*20, 21*). A leading tip cell extends filopodia and lamellipodia to explore its surroundings, while trailing stalk cells elongate and stabilize the nascent vessel (*20, 22, 23*). This process generates a highly reproducible vascular pattern, in marked contrast to the network-like arrangements formed by endothelial collectives under less spatially constrained conditions.

Part of this stereotypy is imposed by guidance signals from the surrounding tissues. Somite-associated Sema3A cues help restrict lateral ISV growth, and both *sema3aa* and *sema3ab* have been implicated by knockdown experiments (*24*). PlexinD1 has an established role in ISV patterning, while recent evidence also identifies Nrp1 as a mediator of Sema3A-dependent endothelial repulsion; the precise plexin partner in this response remains unresolved (*25*). The extracellular matrix (ECM) may provide an additional, overlapping constraint. The intersomitic space is enriched in Laminin-111, Laminin-411, and fibronectin (*26, 27*). Combined loss of *lama1* and *lama4* disrupts ordered ISV outgrowth despite relatively normal sprout initiation, suggesting that laminin contributes to the maintenance rather than the initiation of directional sprouting (*28*). Fibronectin is likewise concentrated at somite boundaries (*27*), while integrin-dependent adhesion, ECM remodeling, and mechanotransductive signaling regulate endothelial polarization and persistent migration (*29, 30*). Patterned laminin can also direct collective cell migration in other developmental contexts *in vivo* (*31*). These findings identify laminin and fibronectin as candidate constraints on endothelial self-organization, but whether they cooperate to maintain stereotyped ISV trajectories remains unclear.

We therefore hypothesized that laminin and fibronectin help channel an intrinsic endothelial capacity for network formation toward stereotyped angiogenic sprouts. This hypothesis predicts that partial weakening of the ECM constraint should reduce the efficiency of directional sprouting, whereas stronger perturbation should permit ectopic interactions between neighboring sprouts and reveal network-like endothelial organization. An important alternative explanation, however, is that laminin and fibronectin perturbation disrupts somite or trunk organization and thereby alters the spatial pattern of established guidance signals such as Semaphorins. Distinguishing a loss of ECM-dependent constraint from secondary disruption of Semaphorin guidance therefore requires examining both vascular and surrounding tissue organization, as well as the expression of relevant Semaphorin genes.

To test this hypothesis, we combined live imaging with perturbation of *lama1*, *lama4*, *fn1a*, and *fn1b*, fibronectin mRNA rescue, analysis of *sema3aa* and *sema3ab* expression, and mathematical modeling. Our model couples an established mechanism of endothelial network formation (*12, 18*) to a deformable ECM (*32, 33, 34*) and Semaphorin-mediated repulsion. This combination allowed us to ask whether changing ECM properties can shift endothelial organization between stereotyped sprouting and lateral fusion into network-like arrangements. Together, the experimental and modeling results support a guided self-organization model (*35, 36*) in which laminin and fibronectin may act alongside other tissue-derived cues to channel endothelial network-forming behavior toward reproducible intersegmental trajectories.

## 2 Results

### 2.1 Laminin and fibronectin support ISV elongation and patterning

To determine how the extracellular matrix (ECM) contributes to intersegmental vessel (ISV) development, we first examined laminin and fibronectin localization during trunk angiogenesis. Using *TgBAC(lama1:lama1-sfGFP)*, *Tg(fn1a:fn1a-mNeonGreen)*, and *Tg(fn1b:fn1b-mCherry)* reporter lines, we observed enrichment of these ECM components in the intersomitic region before and during ISV outgrowth (Figure 1B– D). Endothelial protrusions extended in close proximity to these ECM-rich regions (Videos 1–3). Additional double-transgenic images acquired during primary ISV outgrowth illustrate the spatial relationship between the ECM-reporter signal and extending *kdrl*-positive ISVs (Figure S6).

**Figure 1:**
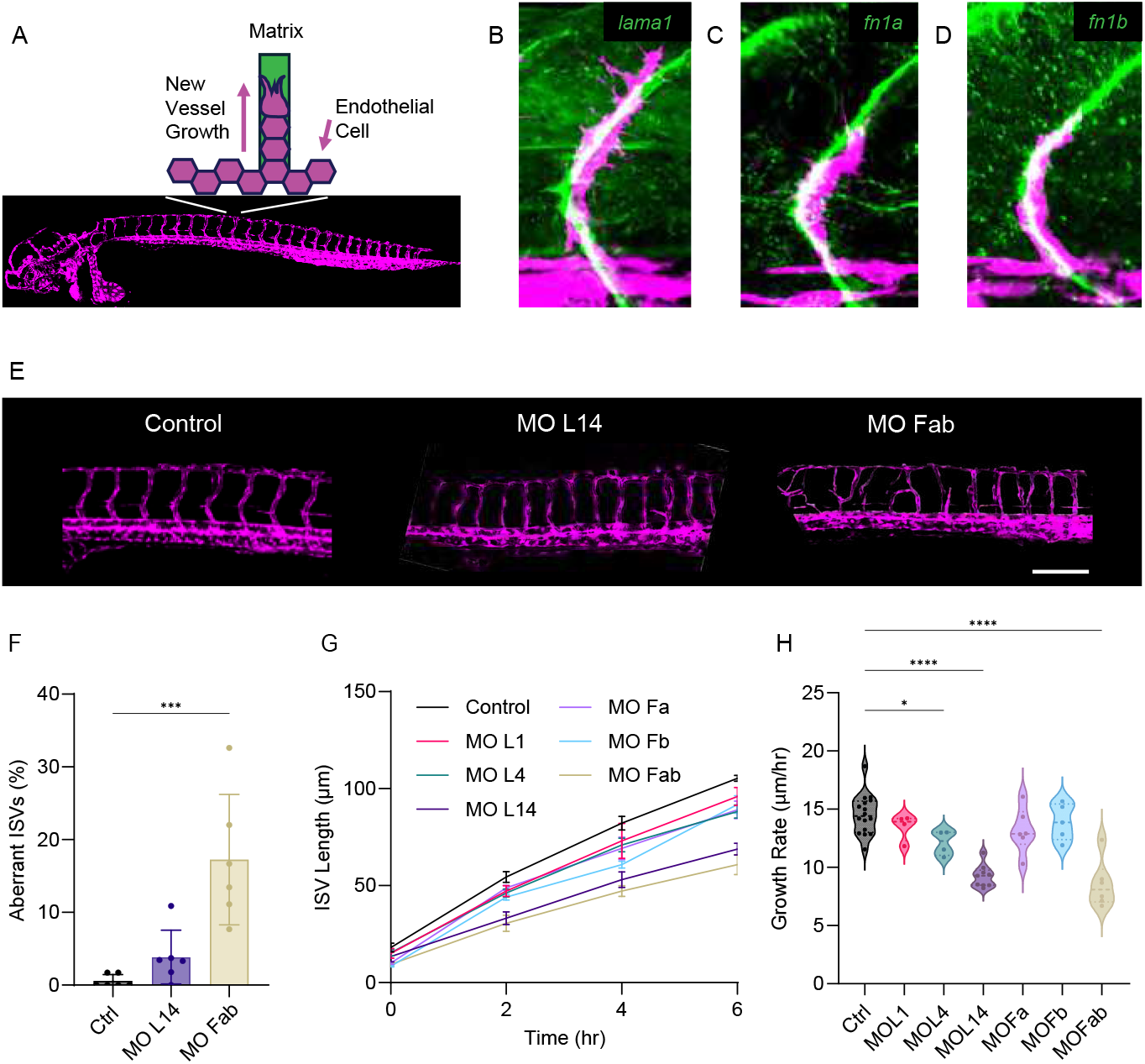
Laminin and fibronectin support efficient ISV elongation and stereotyped patterning. (A) Schematic of ISV sprouting from the dorsal aorta through the intersomitic space. Anterior-left, dorsal-up. (B–D) Representative live images of *TgBAC(lama1:lama1-sfGFP)*, *Tg(fn1a:fn1a-mNeonGreen)*, and *Tg(fn1b:fn1b-mCherry)* reporter signal during ISV development, showing ECM enrichment adjacent to sprouting endothelial cells. (E) Representative higher-magnification confocal images of trunk vasculature in Control, MO L14, and MO Fab embryos at 48 hpf. (F) Percentage of aberrant ISVs per embryo. Individual points represent embryos; bars show mean ± s.d. Kruskal–Wallis test followed by Dunn’s multiple-comparisons test; *n* = 6 embryos per group. (G) Absolute ISV length (*μ*m) during live imaging from 22 to 28 hpf, shown as mean ± s.e.m. (H) ISV growth rate (*μ*m/h) over the 6-h interval. Individual points represent embryos; violin plots show the distributions. One-way ANOVA followed by Dunnett’s multiple-comparisons test; Control *n* = 16, MO L1 *n* = 4, MO L4 *n* = 4, MO L14 *n* = 9, MO Fa *n* = 6, MO Fb *n* = 5, and MO Fab *n* = 6. Statistical significance: \**p* < 0.05, \*\**p* < 0.01, \*\*\**p* < 0.001, \*\*\*\**p* < 0.0001. All embryo images in (B–E) are oriented with anterior to the left and dorsal up. Scale bar, 200 *μ*m.

We initially tested Cas13d-mediated RNA depletion as an orthogonal perturbation strategy. Despite a robust positive-control *tbxta* phenotype, Cas13d targeting of *lama1*, *lama4*, *fn1a*, and *fn1b* did not produce reproducible transcript depletion by RT-qPCR (Figure S1). Cas13d was therefore not used as independent validation of the vascular phenotype; methodological details are provided in the Methods and Supplementary Information. We therefore used dose-titrated morpholino perturbations of *lama1*, *lama4*, *fn1a*, and *fn1b* for the functional experiments. Dose-response testing and reporter-fluorescence measurements are shown in Figure S2; final working doses were selected within the titrated ranges to balance target attenuation with embryo health and are reported in Methods.

At 48 hpf, control embryos showed the stereotyped segmental ISV pattern, whereas combined ECM perturbation produced abnormal vascular trajectories (Figure 1E, F). Quantification clarified the severity of these effects: the percentage of aberrant ISVs per embryo, where ’aberrant’ was defined by ectopic branching, loss of the normal dorsoventral trajectory, or abnormal inter-ISV connections, was significantly increased in MO Fab, whereas MO L14 did not show a significant increase (Figure 1F).

To quantify early ISV extension directly, we performed time-lapse imaging from 22 to 28 hpf and measured absolute ISV length in micrometers. The resulting length trajectories are shown in Figure 1G. ISV growth-rate analysis showed significant reductions in MO L4, MO L14, and MO Fab relative to Control, with the largest reductions in MO L14 and MO Fab; MO L1, MO Fa, and MO Fb did not differ significantly from Control (one-way ANOVA followed by Dunnett’s multiple-comparisons test, Figure 1H). To assess whether differences in embryo dimensions contributed to these results, we repeated the analysis after normalizing the embryo-level mean ISV length to the dorsal– ventral (D/V) axis length of each embryo. The normalized time courses and growth rates showed a similar overall pattern of reduced elongation in the affected ECM knockdown conditions (Figure S4A,B).

Together, these data separate two effects of ECM perturbation: reducing laminin or fibronectin availability can impair the efficiency of early ISV elongation, whereas pronounced patterning defects are not an obligatory consequence at these perturbation levels. This distinction motivated us to ask whether changes in ECM mechanics or attachment-site availability could account for the growth phenotype and under which conditions weaker spatial guidance might additionally promote ISV misguidance.

### 2.2 Reduced ECM stiffness or fiber density can account for reduced ISV growth *in silico*

The experiments above show that laminin/fibronectin perturbation reduces early ISV growth. To ask whether changes in cell–ECM coupling could account for this phenotype, we used a hybrid Cellular Potts model in which elongated endothelial cells co-attract through a secreted chemokine and thereby self-organize into networks (*12, 18*). We extended this network-forming mechanism with a deformable fibrous ECM, mechanosensitive focal adhesions (*33, 32, 34*), and Semaphorin-mediated spatial guidance (Figure 2A).

**Figure 2:**
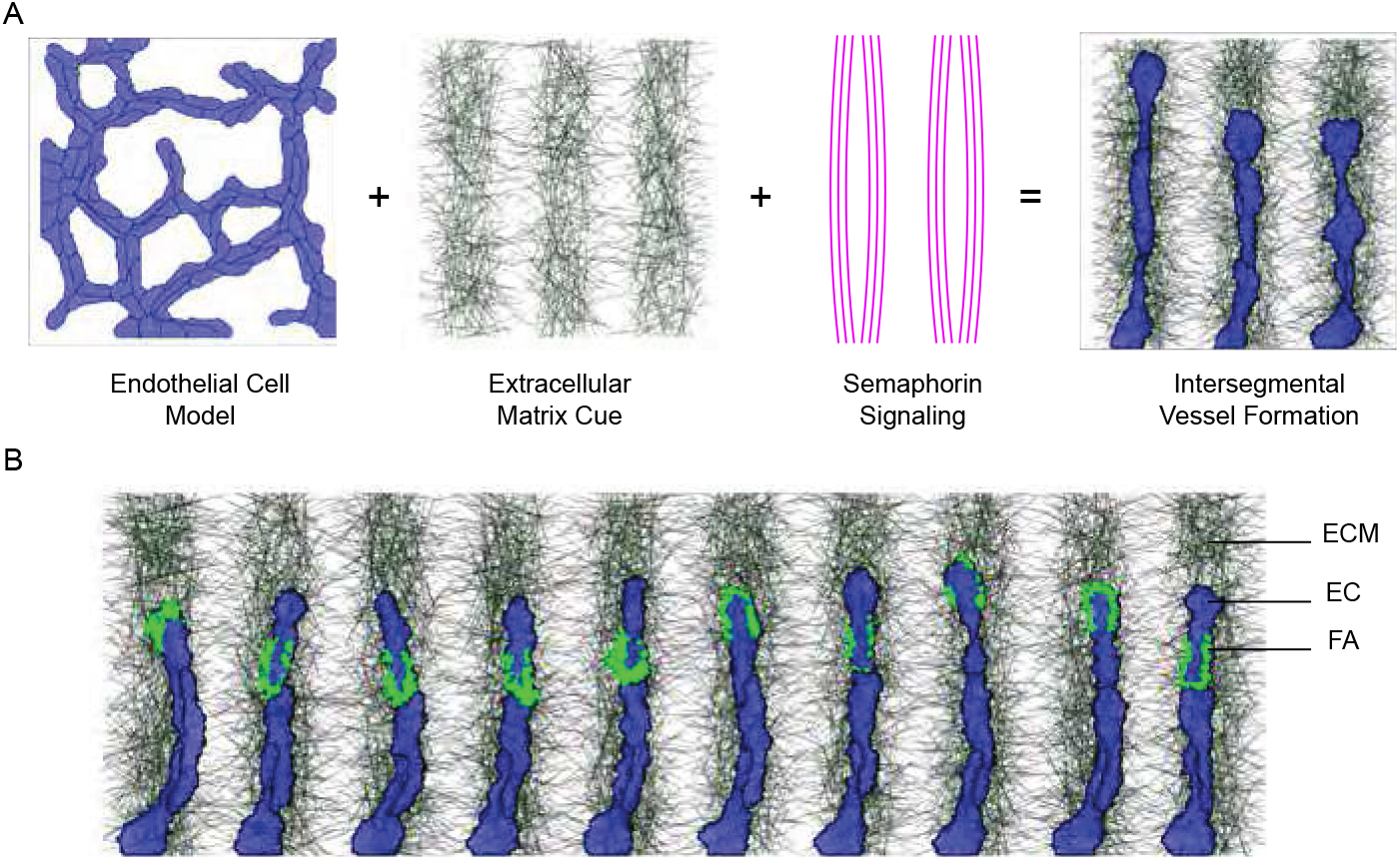
Model overview of ECM-guided ISV sprouting. A: ISV formation in the model arises when intrinsic EC network-forming behavior is constrained by patterned ECM guidance and Semaphorin-mediated repulsion. The panel illustrates the three main subcomponents of the model: EC network formation dynamics, a mechanical model of the ECM, and a model describing semaphorin-mediated chemical signaling. B: Simulation snapshot showing labeled ECs, FAs, and ECM.

Each simulated ISV consisted of one endothelial tip cell followed by two stalk cells (Figure 2A). Based on reported focal-adhesion localization in endothelial tip-cell filopodia (*29*), the baseline model permitted mechanosensitive focal adhesions only in the tip cell, where their formation was polarized toward the leading edge; stalk cells did not form focal adhesions. A generalized fibrous ECM and short-ranged Semaphorin repulsion provided mechanical and spatial guidance; full implementation details are provided in Methods.

We considered two non-exclusive ways in which laminin/fibronectin depletion could alter cell–ECM coupling: reduced ECM mechanical resistance (hypothesis I) or reduced availability of cell–ECM attachment sites (hypothesis II). Because neither property was measured directly in the morphants, we treated reduced stiffness and reduced attachment-site density as alternative mechanistic hypotheses rather than literal representations of the knockdown conditions.

Reducing either ECM stiffness or fiber density significantly reduced simulated ISV growth rate while still allowing sprouts to form, whereas final vessel length did not differ significantly among conditions (Figure 3A–D). The perturbations produced distinct focal-adhesion dynamics: both reduced the total number of FAs, with a stronger reduction under decreased stiffness than under decreased density (Figure 3E), and both shortened FA lifetime, again more strongly under decreased stiffness (Figure 3F). Thus, within the model, reduced stiffness and reduced attachment-site availability weaken effective cell–ECM coupling through partly different effects on adhesion formation and persistence, but converge on reduced sprout extension. Allowing stalk cells to form a spatially unpolarized distribution of FAs reduced growth and eliminated sensitivity to ECM stiffness (Figure S7). This control tested whether stiffness sensitivity depended on polarized tip-cell adhesion; subsequent simulations therefore used the polarized tip-cell FA configuration.

**Figure 3:**
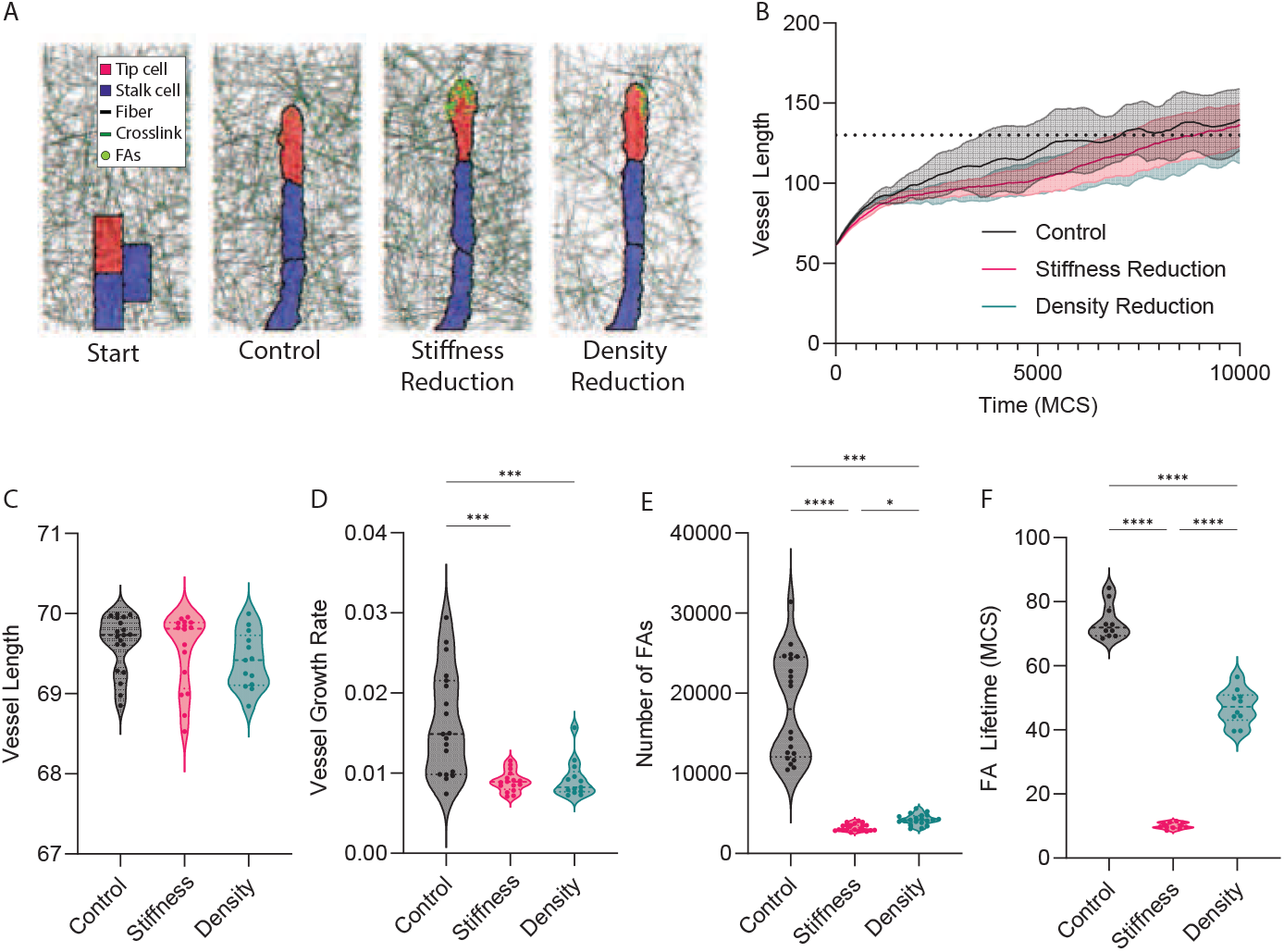
Reduced ECM stiffness or density slows ISV growth in silico. (A) Representative simulation snapshots showing vessel formation. From left to right: initial configuration, Control with stiff and dense ECM, reduced fiber stiffness, and reduced fiber density. (B) Vessel length over simulation time. Lines show the mean and shaded regions indicate ± s.d.; the dotted horizontal line indicates the vessel-length reference used for growth-rate measurement. (C) Final vessel length. No significant differences were detected among conditions (Kruskal–Wallis test followed by Dunn’s multiple-comparisons test; all adjusted *p* > 0.05). (D) Vessel growth rate. Growth rate was reduced under both stiffness reduction (Control vs. Stiffness, adjusted *p* = 0.0004) and density reduction (Control vs. Density, adjusted *p* = 0.0008), whereas Stiffness and Density did not differ significantly (adjusted *p* > 0.9999; Kruskal–Wallis test followed by Dunn’s multiple-comparisons test). (E) Total number of focal adhesions (FAs) formed during each simulation. FA number differed between Control and Stiffness (*p* < 0.0001), Control and Density (*p* = 0.0002), and Stiffness and Density (*p* = 0.0101; Kruskal–Wallis test followed by Dunn’s multiple-comparisons test). (F) Mean FA lifetime per simulation. All pairwise comparisons were significant (ordinary one-way ANOVA followed by Tukey’s multiple-comparisons test; adjusted *p* < 0.0001 for all comparisons). For (C–F), violin plots show the distributions and individual points represent independent simulations. For (C,D), Control *n* = 19, Stiffness *n* = 16, and Density *n* = 13; for (E), *n* = 20 simulations per condition; and for (F), *n* = 10 simulations per condition. Statistical significance: \**p* < 0.05, \*\*\**p* < 0.001, \*\*\*\**p* < 0.0001.

### 2.3 Reduced ECM density or stiffness promotes lateral ISV fusion ***in silico***

We next extended the model to represent ten neighboring ISVs to ask how ECM and Semaphorin guidance influence interactions between adjacent sprouts (Figure 2B). Intersegmental spaces were represented as ECM-rich regions bordered by Semaphorin-mediated repulsive guidance, consistent with the spatial organization of these cues in the zebrafish trunk (*24*) and with our ECM reporter observations (Figure 1B–D).

We varied ECM density or stiffness together with the range of Semaphorin guidance (Figure 4 and Figure S9). Under the reference condition, neighboring ISVs grew in parallel and remained separated. Reducing ECM density or stiffness, or shortening the range of Semaphorin guidance, increased fusion between neighboring simulated ISVs (Figure 4A–C and Figure S9A–C). Thus, the model suggests that overlapping ECM and Semaphorin cues may jointly constrain lateral interactions between neighboring sprouts, supporting the possibility that the ECM acts as one component of ISV guidance.

**Figure 4:**
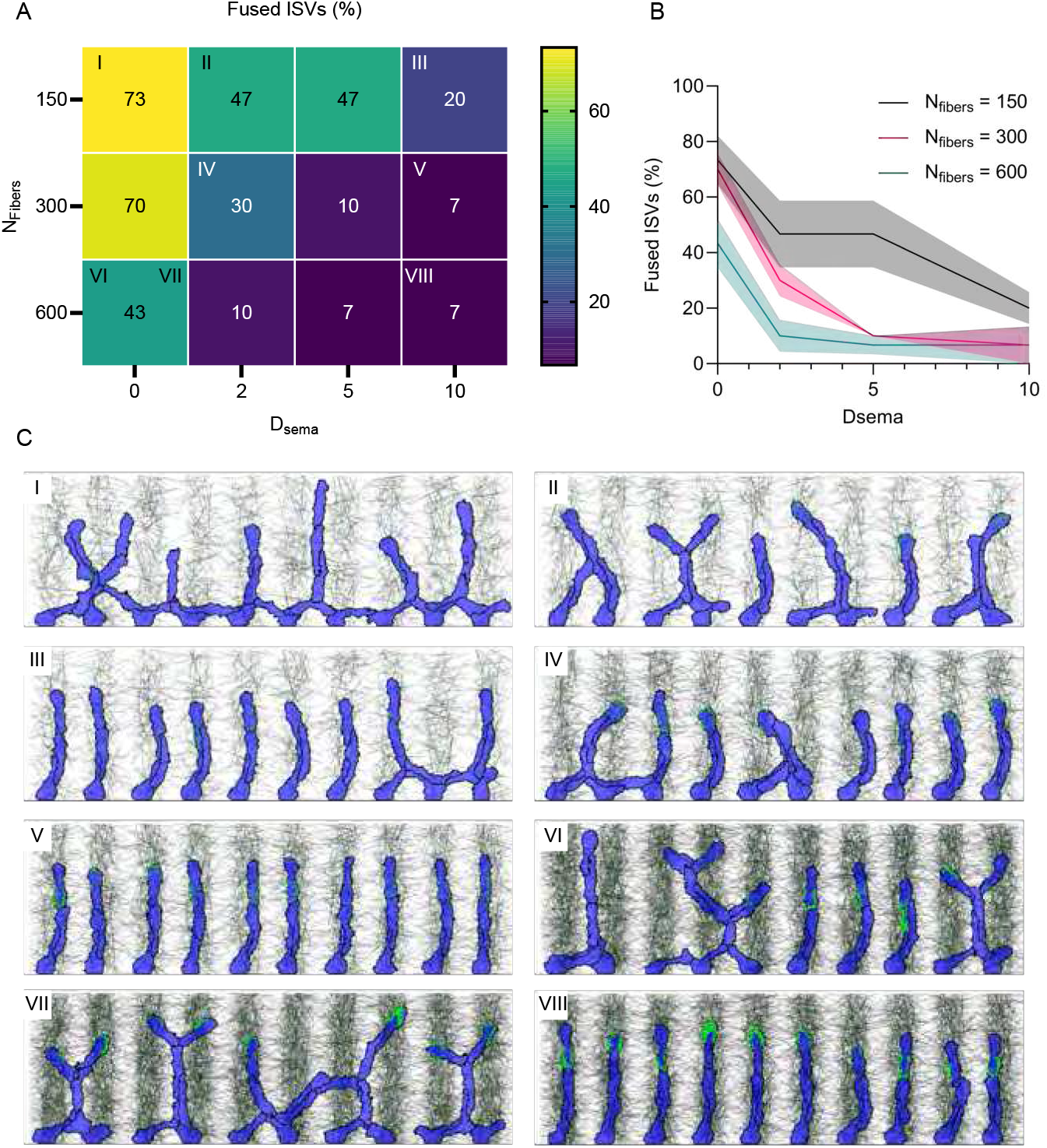
Reduced ECM guidance promotes lateral ISV fusion in silico. Simulations of ISV sprouting as a function of ECM density (*N*_fibers_) and Semaphorin range parameter (*D*_sema_). A: Heatmap showing the percentage of fused vessels for combinations of *N*_fibers_ and *D*_sema_ with fiber stiffness *E*_spring_ = 100 over *n* = 3 independent simulations. Low values indicate well-separated ISVs whereas high values indicate lateral fusion and network formation. B: Percentage of fused vessels as a function of *D*_sema_ and ECM density. Average and standard deviation over *n* = 3 simulations. C: Representative simulation snapshots for cases I-VIII indicated in panel A.

### 2.4 ISV misguidance after combined laminin and fibronectin perturbation

The preceding experiments showed that combined laminin or combined fibronectin perturbation reduced early ISV growth rates, whereas a significant increase in aberrant ISV patterning was detected only after combined fibronectin depletion. We therefore asked whether further reducing ECM redundancy would produce a stronger guidance phenotype. Combining MO L14 with either *fn1a* (MO L14Fa) or *fn1b* (MO L14Fb) significantly increased the percentage of aberrant ISVs (Figure 5A). Absolute ISV-length trajectories from 22 to 28 hpf are shown in Figure 5B, and ISV growth-rate analysis confirmed significantly reduced extension in both combined perturbations relative to Control (Figure 5C). D/V-axis normalization of the embryo-level ISV measurements likewise showed reduced elongation and growth rates in MO L14Fa and MO L14Fb, indicating that differences in embryo dimensions alone cannot account for the observed growth phenotype (Figure S4C,D). Representative trunk images illustrate the associated loss of stereotyped segmental trajectories (Figure 5D). Because combined ECM perturbation can also alter trunk morphology, we asked whether the vascular phenotype might be secondary to broader developmental disruption. Whole-embryo bright-field images showed that gross trunk distortion was more pronounced in MO L14Fa than in MO L14Fb (Figure S5), whereas strong vascular disorganization was present in both conditions. These bright-field observations assess gross tissue morphology only; molecular somite patterning was not measured. In addition to ectopic branching and misrouting, time-lapse imaging directly captured a lateral joining event between adjacent ISVs, confirming that at least a subset of abnormal inter-ISV connections represent vessel fusion (Video 4).

**Figure 5:**
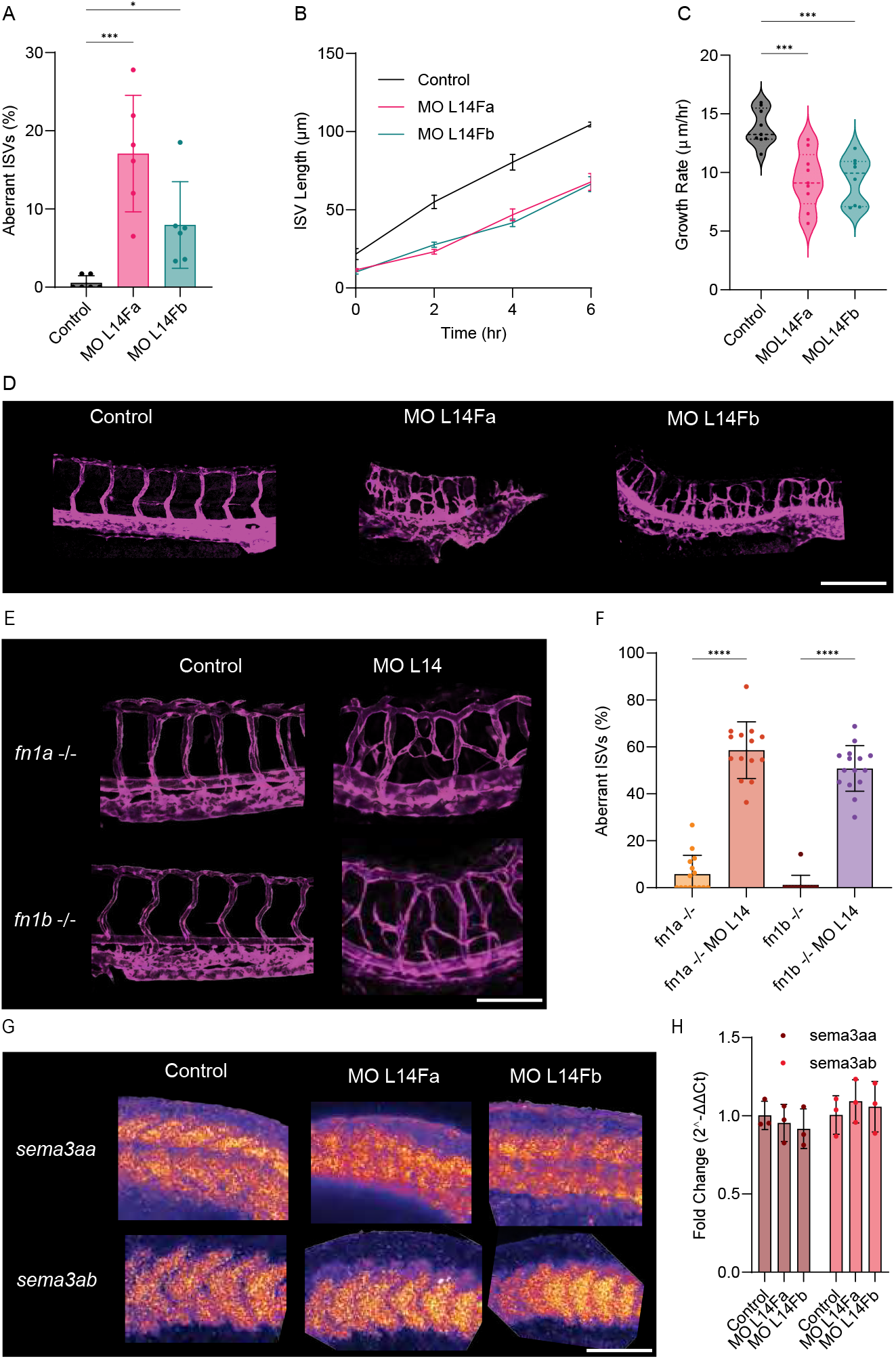
ISV misguidance after combined laminin/fibronectin perturbation and supporting genetic and Semaphorin analyses. (A) Percentage of aberrant ISVs per embryo in Control, MO L14Fa, and MO L14Fb embryos. Individual points represent embryos; bars show mean ± s.d. Kruskal–Wallis test followed by Dunn’s multiple-comparisons test; *n* = 6 embryos per group. (B) Absolute ISV length (*μ*m) during live imaging from 22 to 28 hpf, shown as mean ± s.e.m. (C) ISV growth rate (*μ*m/h) over the 6-h imaging interval. Individual points represent embryos; violin plots show the distributions. One-way ANOVA followed by S^̌^idák’s multiple-comparisons test; Control *n* = 9, MO L14Fa *n* = 9, and MO L14Fb *n* = 8. (D) Representative confocal images of Control, MO L14Fa, and MO L14Fb trunk vasculature. (E) Representative trunk vasculature in homozygous *fn1a^ya^*^13^*^Tg^* and *fn1b^ya^*^14^*^Tg^* mutant embryos, denoted *fn1a*^−/−^ and *fn1b*^−/−^, respectively, with or without MO L14. (F) Percentage of aberrant ISVs in the genetic-validation experiment. Individual points represent embryos; bars show mean ± s.d. Kruskal–Wallis test followed by Dunn’s multiple-comparisons test; *fn1a*^−/−^ *n* = 15, *fn1a*^−/−^ + MO L14 *n* = 14, *fn1b*^−/−^ *n* = 12, and *fn1b*^−/−^ + MO L14 *n* = 15. (G) Representative whole-mount HCR RNA-FISH for *sema3aa* and *sema3ab* in Control, MO L14Fa, and MO L14Fb embryos at 22 hpf. For each Semaphorin target, *n* = 15 embryos were analyzed per morpholino condition. HCR was used for qualitative comparison of spatial expression patterns; fluorescence intensity was not used for quantitative between-group analysis. (H) RT-qPCR analysis of *sema3aa* and *sema3ab*, normalized to *cdk2ap2* and displayed as fold change (2^−ΔΔ^*^Ct^* ) relative to Control. Bars show mean ± s.d.; *n* = 3 biological replicates per condition. Statistical analyses were performed on Δ*Ct* values separately for each gene using ordinary one-way ANOVA followed by Dunnett’s multiple-comparisons test against Control. No significant differences were detected for either *sema3aa* or *sema3ab* (*p* > 0.05). Statistical significance: \**p* < 0.05, \*\**p* < 0.01, \*\*\**p* < 0.001, \*\*\*\**p* < 0.0001. Embryo and trunk images in (D,E,G) are oriented with anterior to the left and dorsal up. Scale bars: 250 *μ*m.

To reduce reliance on morpholino-mediated fibronectin depletion, we next combined MO L14 with genetic loss of *fn1a* or *fn1b*. In both mutant backgrounds, additional laminin depletion strongly increased the percentage of aberrant ISVs relative to the corresponding mutant without MO L14 (Figure 5E,F). These experiments provide orthogonal genetic support for a cooperative laminin–fibronectin contribution to ISV patterning, although the laminin component remains morpholino-based.

To investigate a possible secondary effect of altered trunk or somite organization, we examined the expression of *sema3aa* and *sema3ab*. Qualitative whole-mount HCR RNA-FISH at 22 hpf showed that both transcripts remained detectable in the trunk across conditions. The *sema3ab* transcript pattern appeared broadly preserved, whereas local differences in the *sema3aa* pattern were present in MO L14Fa (Figure 5G). HCR signal intensity varied substantially among specimens and was not used for quantitative between-group comparison. RT-qPCR of whole-embryo RNA at 22 hpf detected no statistically significant differences in either transcript between Control, MO L14Fa, and MO L14Fb, with each gene analyzed separately on Δ*Ct* values (Figure 5H).

Together, these data argue against a wholesale reduction of *sema3aa*/*sema3ab* transcript abundance as the sole explanation for the vascular phenotype. At the same time, the local HCR differences and more pronounced trunk abnormalities in MO L14Fa remain consistent with an additional contribution from altered tissue organization. The strong vascular phenotype in MO L14Fb, despite comparatively preserved gross trunk morphology and a broadly preserved *sema3ab* expression pattern, suggests that secondary tissue disorganization is not the sole explanation. This interpretation is qualitatively consistent with the model result that weakening ECM-dependent guidance increases lateral interactions between neighboring sprouts.

### 2.5 Chimeric fibronectin mRNA rescues abnormal ISV patterning

To test the specificity of the fibronectin contribution to the combined phenotype, we performed chimeric fibronectin mRNA rescue experiments in MO L14Fa and MO L14Fb embryos. The *fn1a* and *fn1b* rescue series were performed as independent experiments in different endothelial reporter backgrounds and were therefore analyzed separately (Figure 6A–C). The chimeric constructs were previously generated and validated for fibronectin morpholino rescue (*27*).

**Figure 6:**
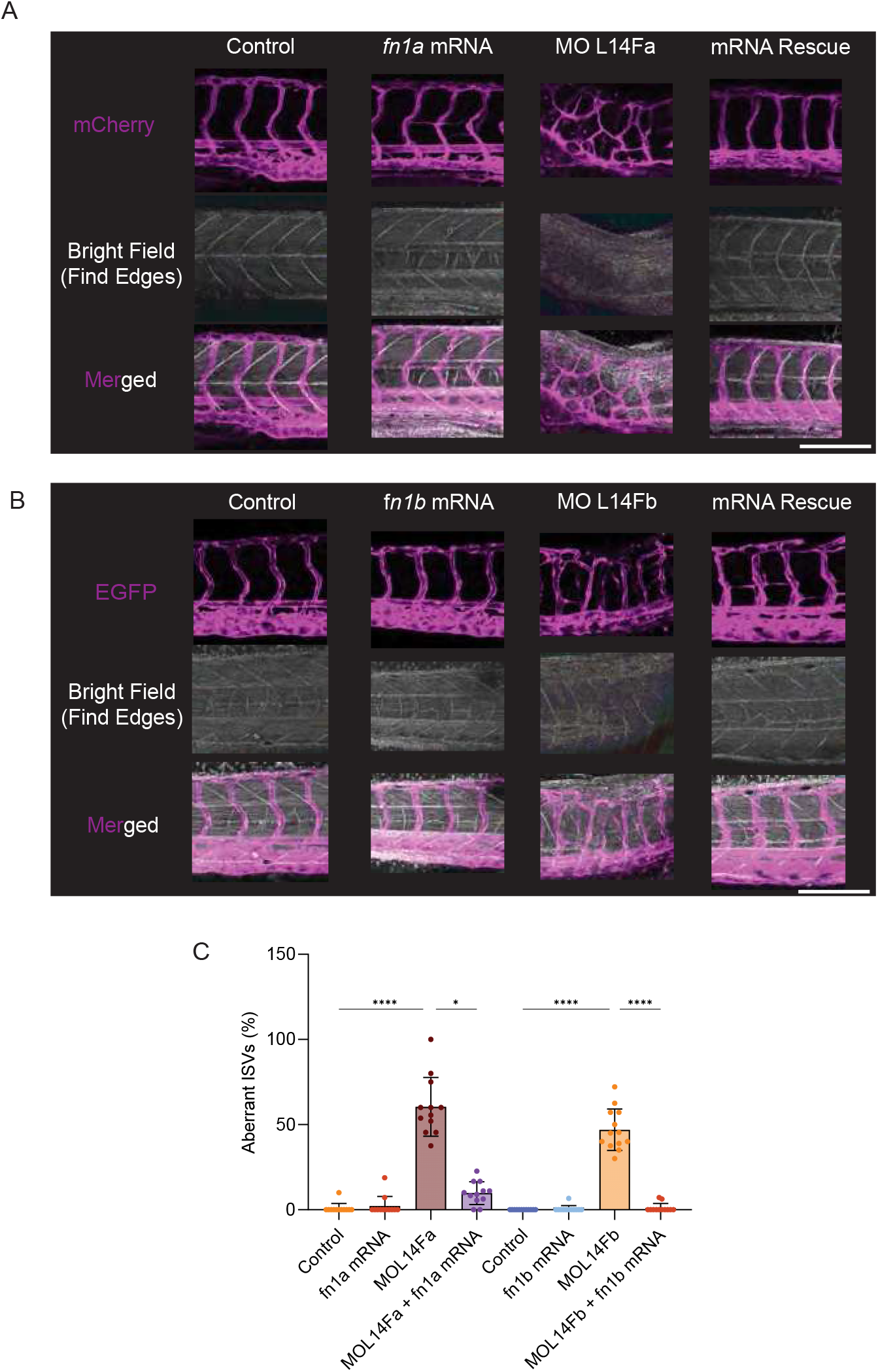
Chimeric fibronectin mRNA rescues abnormal ISV patterning in combined laminin/fibronectin morphants. (A) *fn1a* rescue series in the mCherry endothelial reporter background: Control, *fn1a* mRNA, MO L14Fa, and MO L14Fa + *fn1a* mRNA rescue. For each condition, the endothelial fluorescence channel, the corresponding bright-field image processed with Find Edges in FIJI, and the merged image are shown. (B) *fn1b* rescue series in the EGFP endothelial reporter background: Control, *fn1b* mRNA, MO L14Fb, and MO L14Fb + *fn1b* mRNA rescue. (C) Percentage of aberrant ISVs per embryo in the *fn1a* and *fn1b* rescue series. Individual points represent embryos; bars show mean ± s.d. The two rescue series were performed as independent experiments in different endothelial reporter backgrounds and were analyzed separately by Kruskal–Wallis tests followed by Dunn’s multiple-comparisons tests. For the *fn1a* series, Control *n* = 12, *fn1a* mRNA *n* = 12, MO L14Fa *n* = 12, and MO L14Fa + *fn1a* mRNA *n* = 12. For the *fn1b* series, Control *n* = 12, *fn1b* mRNA *n* = 12, MO L14Fb *n* = 13, and MO L14Fb + *fn1b* mRNA *n* = 12. Aberrant ISVs were defined by ectopic branching, abnormal dorsoventral trajectories, or abnormal inter-ISV connections. Chimeric fibronectin constructs were previously described and validated (*27*). Statistical significance: \**p* < 0.05, \*\**p* < 0.01, \*\*\**p* < 0.001, \*\*\*\**p* < 0.0001. Embryo images in (A,B) are oriented with anterior to the left and dorsal up. Scale bar, 250 *μ*m.

Both MO L14Fa and MO L14Fb showed a strong increase in the percentage of aberrant ISVs relative to their matched controls (Figure 6C). Co-injection of the corresponding chimeric fibronectin mRNA significantly reduced the percentage of aberrant ISVs in both experimental series, supporting rescue of the vascular patterning phenotype. The rescue experiments are interpreted within each matched series rather than by directly comparing the severity of MO L14Fa and MO L14Fb across reporter backgrounds. Representative images show restoration of a more stereotyped segmental vascular pattern after co-injection of the corresponding chimeric fibronectin mRNA in both rescue series (Figure 6A,B). The accompanying bright-field edge images also illustrate that trunk/somite distortion is more pronounced in the MO L14Fa series than in MO L14Fb. Thus, the rescue data support a specific contribution of fibronectin depletion to the combined vascular phenotype while remaining compatible with an additional tissue-organization component, particularly in MO L14Fa.

Together, these chimeric fibronectin mRNA rescue experiments reduce the likelihood that the severe vascular phenotype is explained solely by nonspecific fibronectin morpholino effects and support a cooperative laminin–fibronectin contribution to ISV patterning. Although the whole embryo rescue does not resolve the precise tissue context through which fibronectin restores vascular patterning, the strong vascular phenotype in MO L14Fb despite comparatively preserved gross trunk/somite morphology suggests that the phenotype cannot be explained solely by secondary tissue disorganization; instead, it is consistent with a contribution of laminin/fibronectin to ISV guidance.

## 3 Discussion

In this paper, we asked whether and how the ECM contributes to coordinating development by constraining self-organized cell behavior toward reproducible, stereotyped pattern formation. In particular, we hypothesized that self-organized endothelial network formation and stereotyped ISV growth share the same repertoire of mechanisms, and that knockdown of ECM proteins would ’revert’ ISV formation to endothelial network formation. In particular, we hypothesized that self-organized endothelial network formation and stereotypic growth of ISVs used the same repertoire of mechanism, and hypothesized that knockdown of ECM proteins should ’revert’ ISV formation to endothelial formation. Consistent with this idea, combined knockdown of *lama1*/*lama4* or *fn1a*/*fn1b* reduced early ISV extension, whereas combined laminin–fibronectin perturbation caused pronounced misguidance and abnormal inter-ISV connections. Genetic validation, fibronectin mRNA rescue, and simulations in which reduced ECM fiber density or stiffness slowed sprouting and promoted lateral vessel interactions support a cooperative role for these ECM components. Our results support a model in which the stereotyped geometry of zebrafish ISVs emerges from endothelial network formation constrained by spatial guidance.

An important alternative explanation for the severe vascular phenotype is that combined ECM perturbation alters trunk or somite development. This could disrupt gradients of semaphorins, chemorepellants that repel the ISVs from the somites and are known to play a key role in ISV guidance (*24, 25*). This possibility is particularly relevant to MO L14Fa, which showed pronounced somite abnormalities in addition to ISV misguidance, whereas MO L14Fb embryos retained comparatively preserved gross segmental morphology despite clear vascular disorganization. HCR showed that the *sema3ab* pattern was broadly preserved in both conditions, whereas *sema3aa* showed a more apparent local alteration in the transcript pattern in MO L14Fa despite no significant change in overall transcript abundance. Both *sema3aa* and *sema3ab* have been implicated in ISV patterning (*24*). The broadly preserved *sema3ab* expression pattern, together with continued expression of both paralogues and the RT-qPCR results, argues against a wholesale loss of Semaphorin transcript expression as the sole explanation for the vascular phenotype. Local *sema3aa* alterations and tissue disorganization may nevertheless contribute to the stronger MO L14Fa phenotype. Together with the strong vascular phenotype of MO L14Fb despite comparatively preserved gross segmental morphology, these observations are consistent with an ECM-dependent contribution to ISV guidance that is not readily explained by gross tissue disorganization alone.

The interpretation that ISVs are guided by laminin is consistent with previous evidence that laminin *α*1 and *α*4 have overlapping requirements during ISV development (*28*). Pollard et al. (*28*) showed that simultaneous disruption of laminin *α*1 and *α*4 severely impairs ISV sprouting and extension, with endothelial sprouts failing to migrate correctly along the normally laminin-rich intersegmental regions. Importantly, the same combined, strong perturbation also severely affects notochord differentiation. Thus, while these experiments established an essential role for laminin during ISV development, the experimental conditions did not fully separate direct effects on endothelial guidance from the secondary consequences of general disruption of axial tissues. By contrast, under the approximately 2.7-fold lower, dose-titrated perturbation of laminins that we used, the predominant phenotype was a reduced rate of ISV extension. Severe misguidance became prominent when laminin and fibronectin were perturbed together, consistent with partially overlapping ECM contributions. Differences between the *fn1a* and *fn1b* combinations may additionally reflect their distinct roles in axial tissue organization: *fn1a* is required for normal Fn1b fibrillogenesis in the presomitic mesoderm, whereas *fn1b* is dispensable for normal Fn1a deposition (*27*). These differences provide a plausible basis for the stronger trunk phenotype in MO L14Fa, but do not establish trunk disruption as a sufficient explanation for ISV misguidance.

The genetic-validation and rescue experiments further support the specificity of the combined phenotype. Introducing MO L14 into either *fn1a*^−/−^ or *fn1b*^−/−^ embryos increased aberrant ISVs relative to the corresponding mutant alone, while re-expression of the corresponding chimeric fibronectin mRNA reduced abnormal ISV patterning in both combined morpholino backgrounds. Because the laminin component remains morpholino-based, this represents partial rather than fully genetic validation. Nevertheless, these complementary experiments support a specific cooperative contribution of laminin and fibronectin.

Our reporter fish lines established the presence of Lama1, Fn1a, and Fn1b in the intersomitic region but did not identify the cellular sources of the matrix encountered by sprouting ISVs. Importantly, these ECM components are enriched in the intersomitic region before and during ISV outgrowth (Figure 1B–D and Figure S6), indicating that endothelial sprouts encounter a spatially organized ECM environment as they extend. Published expression data and developmental single-cell transcriptomic datasets identify several candidate source tissues, including notochord, paraxial mesoderm, and muscle (*28,37,27,38*). Pollard et al. further showed that laminin *α*1 can be supplied from non-notochordal tissues (*28*). Endothelial cells can also produce functionally important fibronectin during developmental angiogenesis in other vertebrate systems (*39*), so an endothelial contribution cannot be excluded. More generally, ECM-mediated feedback may warrant future investigation: in vitro evidence (*40*) and computational studies (*41, 42*) suggest that endothelial matrix deposition or remodeling can guide migration along previously modified paths, a form of ’stigmergy’ (*43*). The timing of ECM deposition, together with expression in surrounding axial tissues, is consistent with the idea that the intersomitic matrix is present before ISV outgrowth and contributes to ISV guidance. Laminin and fibronectin perturbation may influence sprouting through several mechanisms, including altered adhesion, ECM architecture and mechanics, tissue organization, and receptor signaling. Targeting individual components of the focal adhesions, including talin or integrin *β*1, which primarily affects endothelial adhesion and migration (*29, 44, 45*), may give more insight into the mechanisms underlying ECM-guided ISV sprouting. In particular, changes in FA lifetime may help distinguish between the alternative mechanisms suggested by the simulations: for reduced ECM stiffness our model predicts that FA lifetime is much more strongly reduced than for reduced ECM density (Figure 3). A further mechanism may involve Dll4/Notch-dependent tip-stalk cell organization, because ECM composition and laminin-binding integrins can modulate Dll4/Notch activity (*46, 47, 48, 49*). Future work should therefore measure Dll4/Notch activity and test whether ECM perturbation alters endothelial cell mixing, including tip-cell overtaking, which has been documented during angiogenic sprouting (*7, 6*).

Several limitations remain. We did not measure VEGF-A expression or its spatial presentation, ECM stiffness or architecture, or the cellular sources of the intersomitic matrix in the morphants. Fibronectin can bind VEGF and modulate endothelial VEGF signaling (*50*), making altered VEGF-A presentation a plausible additional mechanism rather than an established effect of our knockdowns. The model is also two-dimensional and represents the ECM as a generic fibrous network instead of considering the structural differences between laminin and fibronectin. Future experimental work would ideally assess ECM stiffness and organization in combination with tissue-specific perturbations, even if this is technically challenging.

Taken together, our results support a role for laminin and fibronectin in both efficient ISV extension and the maintenance of stereotyped segmental trajectories. Rather than constituting a single dominant guidance system, the intersomitic ECM appears to cooperate with tissue architecture and molecular cues such as Semaphorins to constrain endothelial network-forming behavior. We therefore interpret ISV formation within a guided self-organization framework: intrinsic endothelial collective behaviors contribute to vessel formation, while overlapping environmental cues restrict these behaviors toward reproducible angiogenic trajectories. In this view, self-organization does not replace positional information; rather, it describes endothelial collective behaviors that tissue-derived constraints channel into stereotyped vascular architecture.

## 4 Materials and Methods

### 4.1 Zebrafish maintenance and developmental staging

Leiden University researchers follow European regulations for the use of animals. Wild-type ABTL zebrafish and transgenic strains were maintained and bred under standard conditions (*51*). Lines used in this study included wild-type ABTL, *Tg(kdrl:mCherry-CAAX)* (*52*), *Tg(kdrl:EGFP)* (*53*), *TgBAC(lama1:lama1-sfGFP)* (*31*), *Tg(fn1a:fn1a-mNeonGreen)*, and *Tg(fn1b:fn1b-mCherry)* (*27*), together with the *fn1a^ya^*^13^*^Tg^* and *fn1b^ya^*^14^*^Tg^* mutant alleles used for the genetic-validation experiments (*54*). Mutant embryos were obtained from heterozygous incrosses. Candidate homozygous mutant embryos were initially selected based on the previously described mutant phenotype, and genotype was subsequently confirmed by PCR. Genotyping primer sequences are provided in Supplementary Table 1. One-cell-stage embryos were injected as described below. Developmental age is reported in hours post-fertilization (hpf) for quantitative experiments; embryos were screened for gross morphology under a bright-field stereomicroscope before imaging.

### 4.2 Morpholinos

Antisense MOs (Gene Tools, LLC) targeting 5^′^ sequences near the translation start sites of *lama1*, *lama4*, *fn1a*, and *fn1b* transcripts (Supplementary Table 1) were used in this study. A randomized 25-mer control morpholino was used as the negative control. Working doses were selected by prior titration to balance target attenuation with embryo health (Figure S2); final injected masses and combinations are reported in Section 4.6.

### 4.3 CRISPR-Cas13d knockdown experiments and fibronectin rescue mRNA

Cas13d guide RNAs were designed using the framework of Wessels et al. (*55*) and synthesized by IDT. RfxCas13d/hfCas13d mRNA or purified RfxCas13d protein was co-injected with target-specific guide RNAs at the one-cell stage. A *tbxta* guide-RNA condition served as a positive control and produced the expected phenotype, but targeting of *lama1*, *lama4*, *fn1a*, and *fn1b* did not yield reproducible transcript depletion by RT-qPCR (Figure S1). Cas13d was therefore not used as independent validation of the vascular phenotype. Plasmid construction and recombinant RfxCas13d production followed our previously described workflow (*56*); guide and construct details are listed in Supplementary Table 1.

The pCS2+ chimeric fibronectin rescue constructs were provided by Dörthe Jülich and Scott A. Holley and correspond to constructs previously used for fibronectin rescue (*27*). Plasmids were linearized with NotI-HF and capped mRNA was synthesized using the mMESSAGE mMACHINE SP6 Transcription Kit, purified, quantified, aliquoted, and stored at −80 ^◦^C until use.

### 4.4 RNA purification, cDNA synthesis and RT-qPCR

For each biological replicate, 20 embryos were collected at 22 hpf and total RNA was isolated using TRIzol reagent. RNA concentration was measured using a NanoDrop 2000 and cDNA was synthesized from 1 *μ*g total RNA using the iScript cDNA Synthesis Kit (Bio-Rad). qPCR was performed on a CFX96 Touch system using SsoAdvanced Universal SYBR Green Supermix (Bio-Rad). Primer sequences for *lama1*, *lama4*, *fn1a*, *fn1b*, *sema3aa*, *sema3ab*, and the reference gene *cdk2ap2* are listed in Supplementary Table 1. Relative expression in Figure 5H was calculated using the 2^−ΔΔ^*^Ct^* method with *cdk2ap2* as the reference gene and the matched Control group as calibrator. The experiments comprised three biological replicates per condition.

### 4.5 Whole-mount HCR RNA-FISH

Whole-mount HCR RNA-FISH v3.0 was used to assess the spatial expression of *sema3aa* and *sema3ab*, following the Molecular Instruments whole-mount zebrafish HCR v3 protocol and the principles described by Choi et al. (*57*). Custom probe pools were designed using Pybridizer (*58*) and synthesized by Integrated DNA Technologies (IDT); probe sequences are provided in Supplementary Table 1. Both probe pools used the B1 initiator and were detected with Alexa Fluor 647-conjugated B1 H1/H2 hairpins. Because both targets used the same amplifier channel, *sema3aa* and *sema3ab* were stained in separate embryo samples.

Embryos were collected at 22 hpf, fixed in 4% paraformaldehyde for 1 h at room temperature, dehydrated with methanol and stored at -20 ^◦^C, and subsequently processed according to the manufacturer’s protocol. Probe hybridization was performed overnight at 37 ^◦^C, followed by overnight amplification at room temperature with 2 pmol of each H1/H2 hairpin. Samples were washed in 5×SSCT and mounted in low-melting-point agarose for confocal imaging.

For each target, Control, MO L14Fa, and MO L14Fb samples were processed in parallel and imaged using matched acquisition settings, with *n* = 15 embryos analyzed per morpholino condition for each Semaphorin target. HCR was used for qualitative comparison of spatial expression patterns and not for quantitative between-group fluorescence analysis.

### 4.6 Microinjection, solutions and injection quantities

One-cell-stage embryos were injected with a calibrated 1 nl bolus using a pneumatic microinjector. Morpholino working solutions were dose-titrated before the experiments (Figure S2). For the vascular analyses, the final injected amount was approximately 0.75 ng per embryo for each laminin morpholino (*lama1* and *lama4*), corresponding to a total MO L14 dose of approximately 1.5 ng per embryo, and 5 ng per embryo for each fibronectin morpholino (*fn1a* and *fn1b*). Combined perturbations contained the same dose of each component. The control morpholino was injected at the matched control dose used for the corresponding experiment. For rescue experiments, 600 pg per embryo of the corresponding chimeric *fn1a* or *fn1b* mRNA was co-injected with MO L14Fa or MO L14Fb, respectively.

CRISPR-Cas13d injections used 150–300 pg Cas13d mRNA or 3 ng Cas13d protein together with 450–900 pg total guide RNA in a 1-nl bolus, as indicated for each experiment.

### 4.7 Microscopy and image analysis

Embryos were cleaned and staged under a bright-field stereomicroscope (Leica M50) and screened for fluorescence using a Leica M205 FA fluorescence stereomicroscope. For confocal imaging, embryos at the indicated hpf were mounted in 0.8% low-melting-point agarose and imaged on a Leica Stellaris 5 confocal microscope using 10×, 20×, or 40× objectives. Acquisition settings were kept constant between groups for fluorescence comparisons.

Z-stacks and time-lapse datasets were analyzed in FIJI/ImageJ (*59*). Time-lapse datasets were acquired from approximately 22 to 28 hpf during primary ISV sprouting. Because ISV sprouting progresses from anterior to posterior, three ISVs per embryo that had only recently initiated dorsal extension from the dorsal aorta at the beginning of the recording were selected and followed throughout the time-lapse. ISVs that had already undergone substantial elongation at the first time point were excluded. Thus, measurements were initiated from vessels at a comparable stage of sprout development rather than from a fixed anatomical position along the anterior-posterior axis. The same selection criteria were applied to all experimental groups.

ISV length was measured manually in micrometers at each time point from the dorsal-aorta origin to the distal sprout tip. The three selected ISVs were averaged to obtain one embryo-level mean at each time point. ISV growth rate was calculated for each embryo as the change in mean ISV length between 22 and 28 hpf divided by 6 h and is reported in *μ*m/h. Absolute ISV length and growth rate are used as the primary readouts. As a complementary size-normalized analysis, the embryo-level mean ISV length at each time point was divided by the D/V-axis length of the corresponding embryo, yielding dimensionless normalized length. Normalized growth rate was calculated as the difference between normalized lengths at 28 and 22 hpf divided by 6 h and is reported in h^−1^ (Figure S4).

For patterning analysis, scorable trunk ISVs within the predefined imaging region were classified as normal or aberrant. Aberrant ISVs were defined by ectopic branching, loss of the normal dorsoventral trajectory, or abnormal lateral/inter-ISV connections. The percentage of aberrant ISVs was calculated for each embryo as 100×(aberrant ISVs/total scored ISVs). The embryo was treated as the biological replicate. The term “fusion” was reserved for events in which vessel joining was directly demonstrated by time-lapse imaging; an example is provided in Video 4.

ECM reporter fluorescence was quantified in FIJI from matched confocal acquisitions. Sum-intensity projections were generated and a trunk ROI spanning five consecutive somites was analyzed. Corrected total fluorescence was calculated as CTF = IntDen(ROI) − Area × Mean(BG), using the mean intensity of three background ROIs. All imaging was monitored and visualized in LAS X software (Leica Microsystems) during acquisition.

### 4.8 Statistical analysis

Statistical analyses were performed in GraphPad Prism. The embryo was treated as the biological replicate for all *in vivo* vascular measurements. Figure 1F was analyzed by Kruskal–Wallis test followed by Dunn’s multiple-comparisons test; Figure 1H by one-way ANOVA followed by Dunnett’s multiple-comparisons test; Figure 5A by Kruskal–Wallis test followed by Dunn’s multiple-comparisons test; Figure 5C by oneway ANOVA followed by S^̌^idák’s multiple-comparisons test; and Figure 5F by Kruskal– Wallis test followed by Dunn’s multiple-comparisons test. For the Semaphorin RT-qPCR dataset (Figure 5H), Δ*Ct* values were analyzed separately for *sema3aa* and *sema3ab* using ordinary one-way ANOVA followed by Dunnett’s multiple-comparisons test against Control. The *fn1a* and *fn1b* rescue experiments in Figure 6C were performed as independent experimental series in different endothelial reporter backgrounds and were therefore analyzed separately. Each series was analyzed by Kruskal–Wallis test followed by Dunn’s multiple-comparisons test.

For the supplementary validation analyses, the Cas13d RT-qPCR datasets in Figure S1 were analyzed by Kruskal–Wallis tests, with Dunn’s multiple-comparisons tests against Control where applicable. In Figure S2, panel B was analyzed by t-test. For the CTF dose-titration analyses in panel C, MO L14 and MO Fb were analyzed by ordinary oneway ANOVA followed by Dunnett’s multiple-comparisons test against Control, whereas MO Fa was analyzed by Kruskal–Wallis test followed by Dunn’s multiple-comparisons test.

For the simulation analyses in Figure 3, each independent simulation was treated as the replicate. Final vessel length (Figure 3C), vessel growth rate (Figure 3D), and total FA number (Figure 3E) were analyzed using Kruskal–Wallis tests followed by Dunn’s multiple-comparisons tests. Mean FA lifetime was first calculated separately for each independent simulation, and these simulation-level means were compared by ordinary one-way ANOVA followed by Tukey’s multiple-comparisons test (Figure 3F). The vessel-length trajectories in Figure 3B are shown as mean ± s.d.; growth rate in panel D was used for inferential comparison between conditions.

The ISV-length time courses in Figure 1G and Figure 5B are shown to visualize growth trajectories; embryo-level ISV growth rate over the 6-h interval is used as the primary inferential summary in Figure 1H and Figure 5C. Figure S4 shows the corresponding D/V-normalized time courses and embryo-level growth rates as a complementary analysis. Figure legends report the statistical test, multiple-comparisons procedure, sample size, and graphical summary. Exact adjusted *p* values are retained in the Prism/source-data output.

### 4.9 Mathematical Model

The mathematical model was implemented as a hybrid cellular Potts model (CPM) (*60*) that combines a CPM to model endothelial cells, with partial-differential equation-based simulations of chemotactic gradients (*12*) and molecular dynamics simulations of the ECM (*33*). The CPM is a cell-based modeling technique in which cells are represented by multiple lattice sites. The model follows our previous CPM of endothelial network formation, in which each EC is elongated and secretes a diffusing chemical to which other ECs are attracted. This cell elongation model (*12*) reproduces the temporal dynamics of *in vitro* assays (*18*). In this work, we extend the cell elongation model with a molecular dynamics model describing the mechanics of the ECM (*33*), a model of the FA-mediated cell-ECM interaction (*32, 34*) and a partial-differential model of semaphorin signaling and chemotaxis. The following sections describe each model component.

#### 4.9.1 Cellular Potts model

The CPM represents cells as collections of points on a rectangular lattice Λ of size *L_x_* × *L_y_*. Each lattice site *p* ∈ Λ is assigned a spin *σ_t_* ( *p*) ∈ **Z** where *t* denotes the time, and the spin value uniquely identifies a biological cell, such that a cell is identified as the collection of lattice sites with identical and non-zero spins. The lattice sites with spin equal to 0 describe space which is not occupied by the cell. Cellular dynamics are described by stochastic cell extensions and retractions due to activity of the cytoskeleton. This process is mimicked by Metropolis-Hastings dynamics: Iteratively, two neighboring lattice sites *p* and *p*^′^ at a cell-cell or cell-ECM interface (i.e., *σ_t_* ( *p*) ≠ *σ_t_* ( *p*^′^)) are sampled and the copying of *σ_t_* ( *p*^′^) into *p* is considered. The copy-attempt is then accepted with probability min{1, exp(−Δ*H*/*μ*)}, where *μ* is a motility parameter and where Δ*H* is the energy difference gained by this copying,

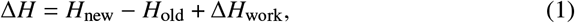

where *H*_old_ and *H*_new_ are the energies in the system before and after the copy-attempt respectively, and Δ*H*_work_ describes an additional, dissipative term describing interactions with the ECM and chemical fields. Time runs in Monte Carlo steps (MCS), i.e., the number of spin interfaces in the grid.

The Hamiltonian energy of the system is defined as

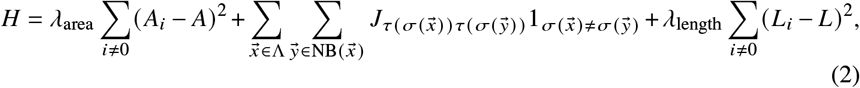

where the first term is an area constraint with coefficient *λ*_area_, *A_i_* is the area of the cell with spin *i*, and *A* is the target area. The second term is a penalty for cell-cell and cell-medium interfaces, with *J_τ_*_,*τ*_′ the penalty size for two neighboring cells of type *τ* and *τ*^′^. The final term is the length constraint with *λ*_length_ a parameter, *L_i_* the length of the cell with spin *i* and *L* is the target length. The lengths of the cells are approximated dynamically during the simulation with the inertia tensor (*12*). The term Δ*H*_work_ in Equation (1) describes any active work generated by the cells due to interactions with the ECM and the chemical field,

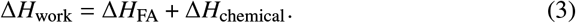

The term Δ*H*_FA_ penalizes retraction from FAs that are potentially present in *p*, and the term Δ*H*_chemical_ gives the cells’ responses to chemical cues, as detailed in the next sections.

#### 4.9.2 Extracellular matrix

The ECM is represented by a mass-spring network of cross-linked identical fibers as described in earlier work (*33, 34*). We briefly summarize the ECM model here and refer to earlier work for a more detailed description of the network construction and mechanics (*33, 34*).

The ECM is modeled as a mass-spring system consisting of beads *x*→_1_, *x*→_2_, . . . , *x*→*_N_* ∈ **R**^2^, connected by harmonic springs with spring constant *E*_spring_. Beads at the boundary of the domain are fixed in place, as are the adhesion beads to which cells attach; the remaining bead positions are updated using an overdamped Langevin equation. Before the integration of the Langevin equation, the cells contract the adhesion beads towards their centers of mass, deforming the ECM locally. By integrating the Langevin equation, these deformations propagate throughout the mass-spring system. The Langevin equation is integrated to steady state after each MCS. We will now discuss how cells react to the ECM by describing the term Δ*H*_FA_ in Equation (3).

Cells interact with the ECM not only by applying contractile forces but also by resisting retractions from adhesion sites through the term Δ*H*_FA_, which penalizes focal adhesion detachment. Each adhesion bead represents a focal adhesion, and the retraction penalty depends on the number of bound integrins:

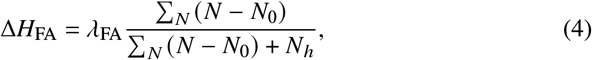

where the sum is over focal adhesions at the retracting site *p*, with *N* the number of bound integrins, *λ*_FA_ a scaling parameter, *N*_0_ the minimum integrin count per adhesion, and *N*_ℎ_ a saturation parameter. The number of bound integrins is updated after each timestep of the CPM and it depends on the tension in the corresponding adhesion bead, as individual focal adhesions behave as catch bonds (*61, 34, 32*). For the exact parameters and implementation of the adhesion particles we refer to (*34*).

#### 4.9.3 Cell polarization

Cell polarization was included by endowing each cell with a polarization vector *v*→(*t*) that is given by a weighted sum of past displacements (*62*) _min{_ *_N_*polarity,hist_,*t*−1}_

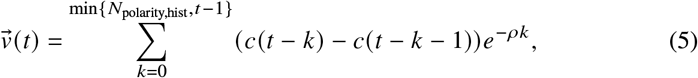

where *c*(*t*) is the cell’s center of mass at time *t* or the zero vector if the simulation run time is less than *t*, *N*_polarity,hist_ is a parameter describing the maximum number of MCS in the history, *ρ* is a decay parameter of the memory effect. Here we have set *N*_polarity,hist_ sufficiently large such that *ρ* determines the cell’s polarization history.

The FA breakdown rate is then determined by the polarization vector, as follows. In the trailing edge of the cell, i.e., at the lattice sites away from the polarized direction, FAs are broken down immediately and removed from the simulation. As a result, FAs can only form at the leading edge of the cell. This effect of the polarization vector on the break-down rate of the FAs at the trailing edge, together with the role of the FAs in stabilizing cellular protrusions leads to a positive feedback loop that drives an ECM-stiffness-dependent persistent random walk (Figure S10 and Videos 5-6) and stiffness-dependent chemotactic sensitivity (Figure S11).

#### 4.9.4 Chemical signaling

To model the gradients of autocrine chemokines that drive EC co-attraction and the semaphorins (*24*), we introduce two chemical fields *c*_chemokine_(*x*→) and *c*_sema_(*x*→).

The semaphorin field, *c*_sema_(*x*→), is assumed static, as

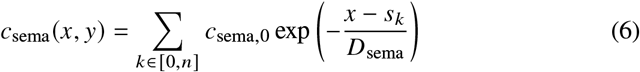

where *D*_sema_ is the characteristic range parameter of the static Semaphorin field, *c*_sema,0_ is the concentration of semaphorins at the somite boundary, and the sum is taken over the somite locations *s_k_*. In simulations with *D*_sema_ = 0, Semaphorin guidance is absent. Somite locations are defined as *s_k_* = *k L_x_*/*n* with *n*, the number of somites and *L_x_*, the width of the simulation domain.

The chemokine signal is assumed to be secreted by the ECs. It is modeled dynamically and diffuses and degrades according to

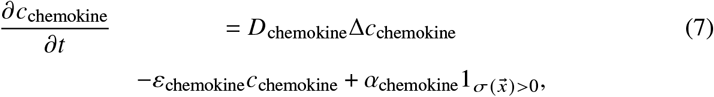

in which *D*_chemokine_ and *ε*_chemokine_ are the diffusion constant and the degradation constant, and *α*_chemokine_ is the secretion rate. After each MCS, Equation (7) is integrated to steady state using a forward Euler scheme.

Chemotaxis is modeled by coupling the CPM through the term Δ*H*_chemical_ in Equation (3). We use the so-called ‘extension-retraction’ chemotaxis algorithm (*13*): Copy-attempts upwards and downwards the signaling gradient add or subtract a small amount of energy to Δ*H* thus biasing the likelihood of acceptance according to the chemical gradient,

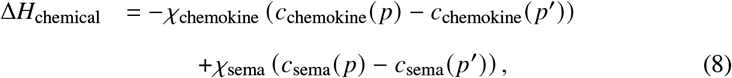

where *χ*_chemokine_ and *χ*_sema_ are parameters describing the sensitivity of the cell to the chemical fields, and negative contributions yield chemoattraction and positive contributions yield chemorepulsion. In the single-cell migration analyses shown in Figures S10 and S11, the imposed directional chemoattractant is denoted VEGF and the corresponding chemotactic sensitivity as *χ*_VEGF_, following the notation used in those simulations; the multicellular ISV model uses *χ*_chemokine_ for EC co-attraction.

#### 4.9.5 Parameter values

The parameter values used in this study are listed in Table 1. All values are dimensionless; robustness of the results was assessed through parameter sensitivity analyses and the parameter variations explored in the simulations.

**Table 1:** Model parameters used in this study.

| Symbol | Description | Value |
| --- | --- | --- |
| **Cellular Potts model** |  |  |
|  | Total simulation time | 10 000 |
| $L_x$ | Domain width for single and multiple ISV simulations | 200, 633 |
| $L_y$ | Domain height | 200 |
| $\mu$ | Cell motility parameter | 50 |
| $A$ | Target cell area | 800 |
| $L$ | Target cell length | 40 |
| $\lambda_{\text{area}}$ | Area constraint coefficient | 1.0 |
| $\lambda_{\text{length}}$ | Length constraint coefficient | 7.0 |
| $J_{\text{cell,cell}}, J_{\text{cell,medium}}$ | Cell–cell and cell-medium adhesion energy | 20, 20 |
| **ECM & focal adhesions** |  |  |
| $N_{\text{fibers}}$ | Number of fibers per ISV | 600 |
| $N_{\text{crosslinks}}$ | Number of crosslinkers per ISV | 2400 |
| $E_{\text{spring}}$ | ECM fiber spring constant (stiffness parameter) | Varied; see figure legends |
| $\rho$ | Polarity update rate | 0.001 |
| $N_{\text{polarity,hist}}$ | Polarity memory length | 100 |
| $N_0$ | Minimal FA size | 50 |
| $N_h$ | Saturation parameter | 70 |
| $\lambda_{\text{FA}}$ | Scaling parameter | 100 |
| **Chemical** |  |  |
| $\chi_{\text{sema}}$ | Chemotactic sensitivity to semaphorin | 1000 |
| $\chi_{\text{chemokine}}$ | Chemotactic sensitivity to chemokine | 1000 |
| $D_{\text{sema}}$ | Semaphorin range parameter | 10 |
| $D_{\text{chemokine}}$ | chemokine diffusion coefficient | $1.0 \times 10^{-13}$ |
| $\varepsilon_{\text{chemokine}}$ | chemokine degradation rate | 0.00018 |
| $\alpha_{\text{chemokine}}$ | Secretion rate | 0.00018 |

## Acknowledgements

We thank Dörthe Jülich and Scott Holley for generously providing the fibronectin fish lines and plasmids, and Caren Norden for providing the laminin fish lines. We are grateful to Vincent Vermeulen for assistance with image analysis and to Rianka M. Eertink for her support during her internship. We are grateful to Herman P. Spaink and Coert Margadant for valuable discussions. We greatly appreciate the excellent support and animal care provided by our fish facility caretakers Guus van der Velden, Michel Mulders, and Ulrike Nehrdich. This publication is part of the ‘Mathematics-based strategies for repairing tumour blood vessels’ project with number 865.17.004 to RM of the Vici research program, which is financed by the Dutch Research Council (https://www.nwo.nl/). The funders had no role in study design, data collection and analysis, decision to publish, or preparation of the manuscript.

## Author notes

Joaqúın Abugattas-Núñez Del Prado and Koen A. E. Keijzer contributed equally to this work. Present address for Erika Tsingos: Theoretical Biology and Bioinformatics, Utrecht University, Utrecht, The Netherlands. Correspondence to.

## Supplementary Figures

**Supplementary Figure 1:**
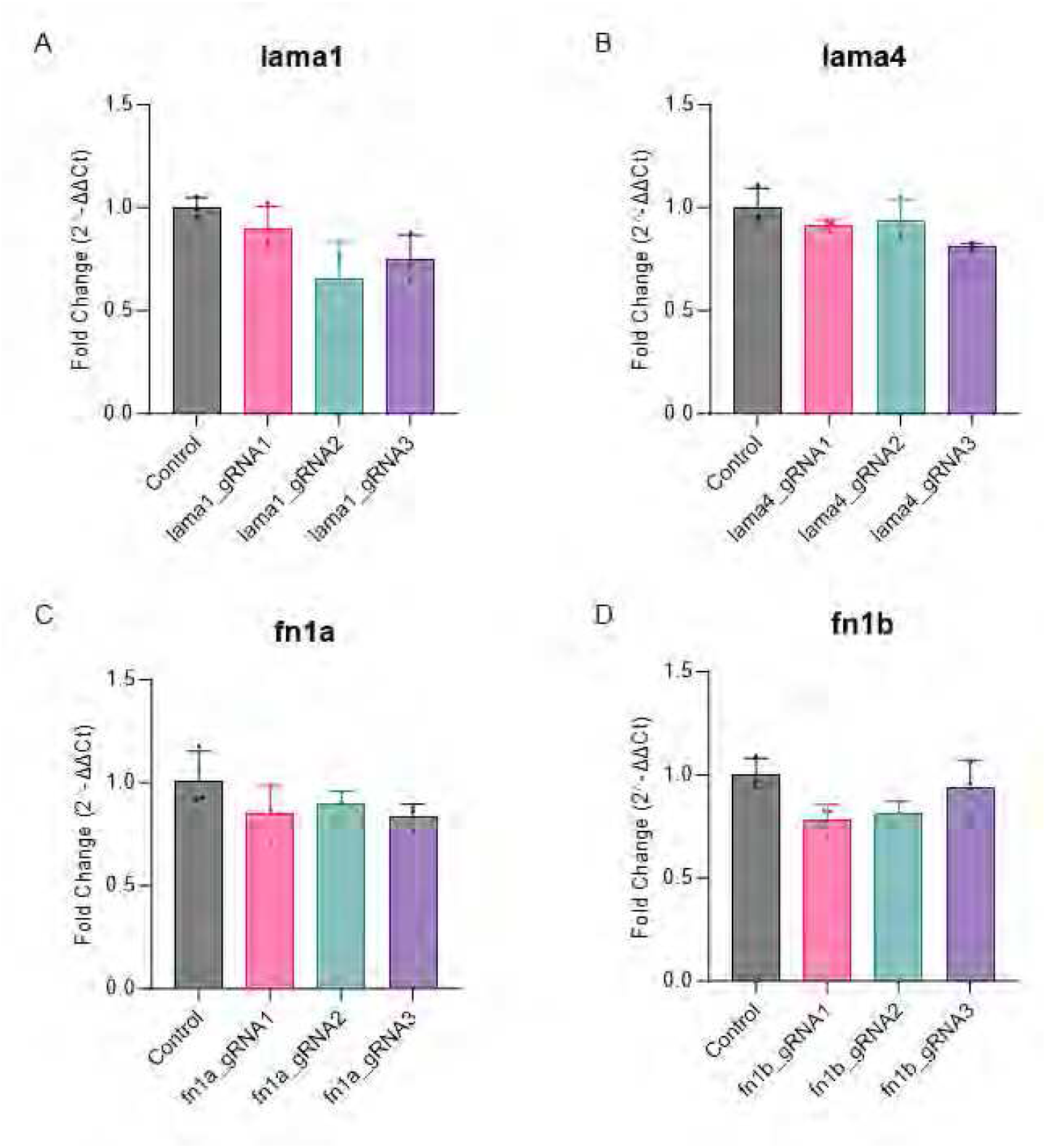
Cas13d-mediated RNA-targeting tests for *lama1*, *lama4*, *fn1a*, and *fn1b*. RT-qPCR analysis of (A) *lama1*, (B) *lama4*, (C) *fn1a*, and (D) *fn1b* transcript abundance in zebrafish embryos injected with three independent gRNAs targeting each transcript, compared with Control. Dots represent biological replicates; *n* = 3 per condition. Cas13d did not produce reproducible transcript depletion across independent gRNAs for any target. Statistical analysis was performed by Kruskal–Wallis test followed by Dunn’s multiple-comparisons test against Control.

**Supplementary Figure 2:**
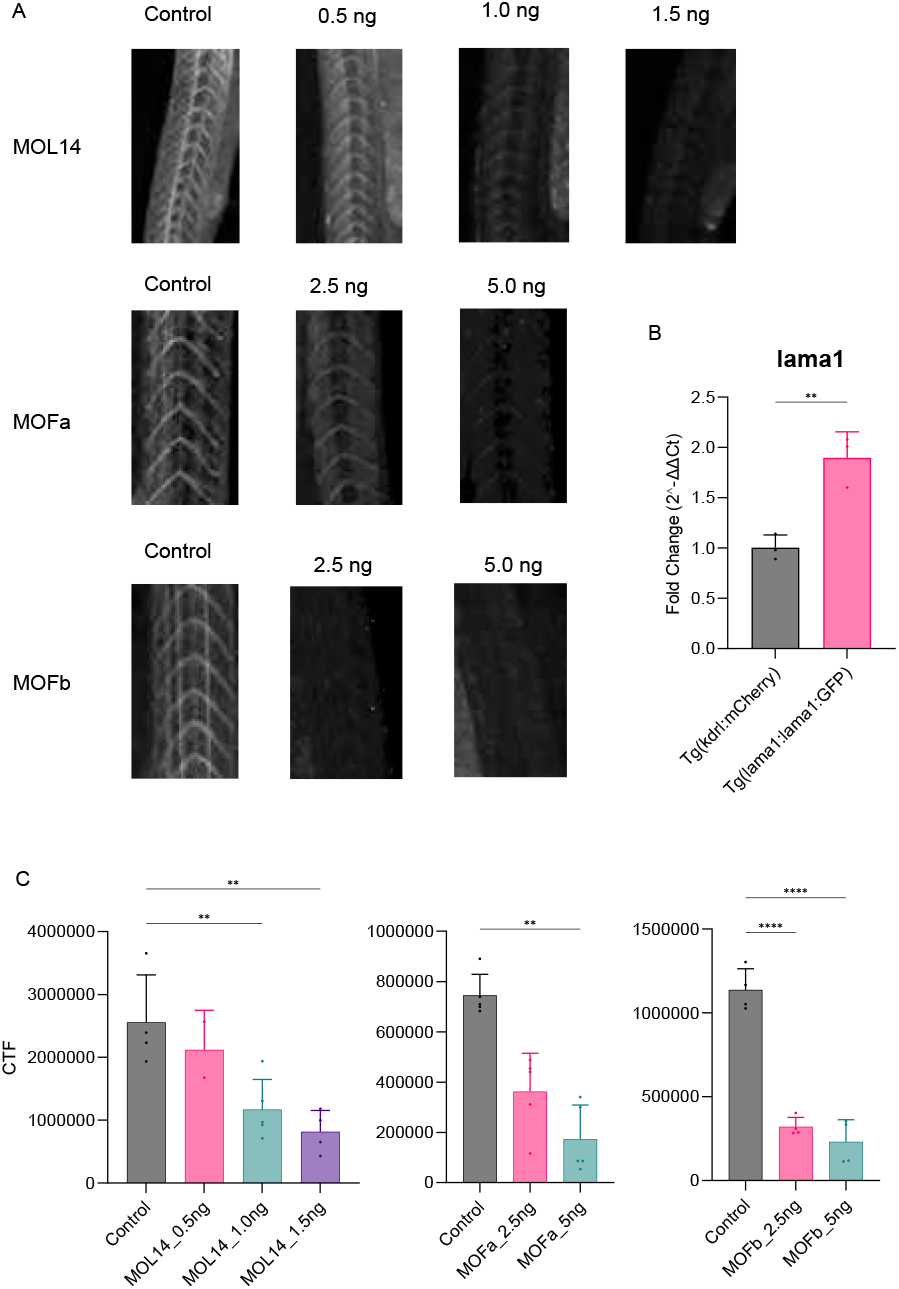
Morpholino dose titration and reporter-fluorescence response. (A) Representative images of zebrafish trunks at 35 hpf for the indicated morpholino (MO) doses. For MO L14, the indicated dose represents the total combined laminin morpholino dose (*lama1* + *lama4*), with equal contribution from each morpholino. (B) Relative *lama1* transcript levels in *Tg(kdrl:mCherry)* and *TgBAC(lama1:lama1-sfGFP)*, analyzed by t-test. (C) Corrected total fluorescence (CTF) across the indicated morpholino titration conditions. MO L14 and MO Fb were analyzed by ordinary one-way ANOVA followed by Dunnett’s multiple-comparisons test against Control; MO Fa was analyzed by Kruskal–Wallis test followed by Dunn’s multiple-comparisons test. For MO L14, Control *n* = 4, 0.5 ng *n* = 2, 1.0 ng *n* = 5, and 1.5 ng *n* = 4; for MO Fa, *n* = 5 per group; for MO Fb, *n* = 4 per group. Statistical significance: \**p* < 0.05, \*\**p* < 0.01, \*\*\**p* < 0.001, \*\*\*\**p* < 0.0001.

**Supplementary Figure 3:**
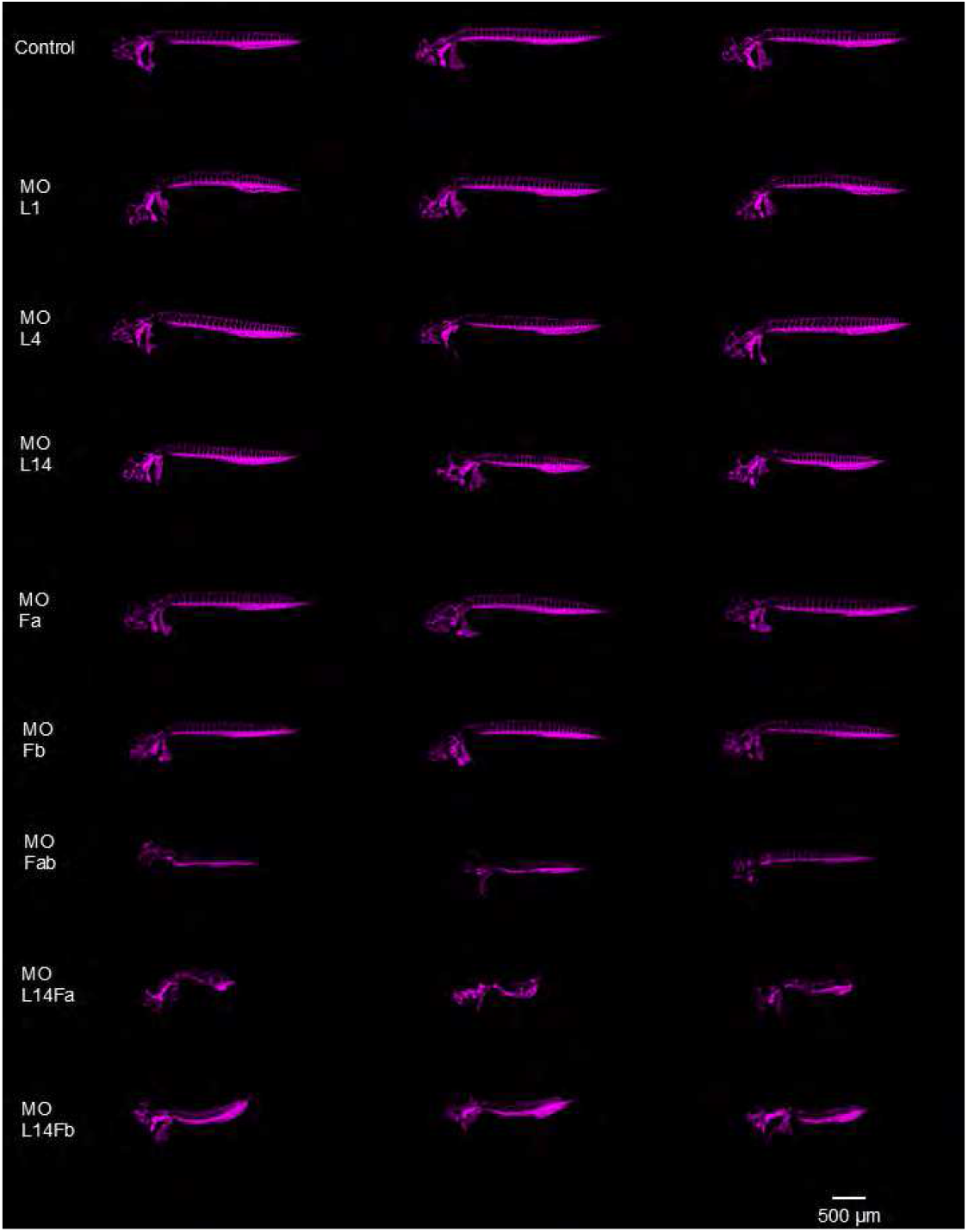
Phenotypic variability across morpholino conditions targeting ECM components. Representative maximum-intensity projections of zebrafish embryos at 48 hpf, showing three examples per condition. Embryos were injected with morpholinos targeting laminin alpha 1 (MO L1), laminin alpha 4 (MO L4), fibronectin 1a (MO Fa), fibronectin 1b (MO Fb), or the indicated combinations (MO L14, MO Fab, MO L14Fa, and MO L14Fb). Control embryos were injected with the randomized 25-mer control MO. Gross trunk abnormalities are most evident in MO L14Fa, whereas MO L14Fb retains comparatively preserved overall trunk morphology despite clear vascular disorganization. The images illustrate phenotypic variability and are not used as a molecular assessment of somite patterning. Scale bar, 500 *μ*m.

**Supplementary Figure 4:**
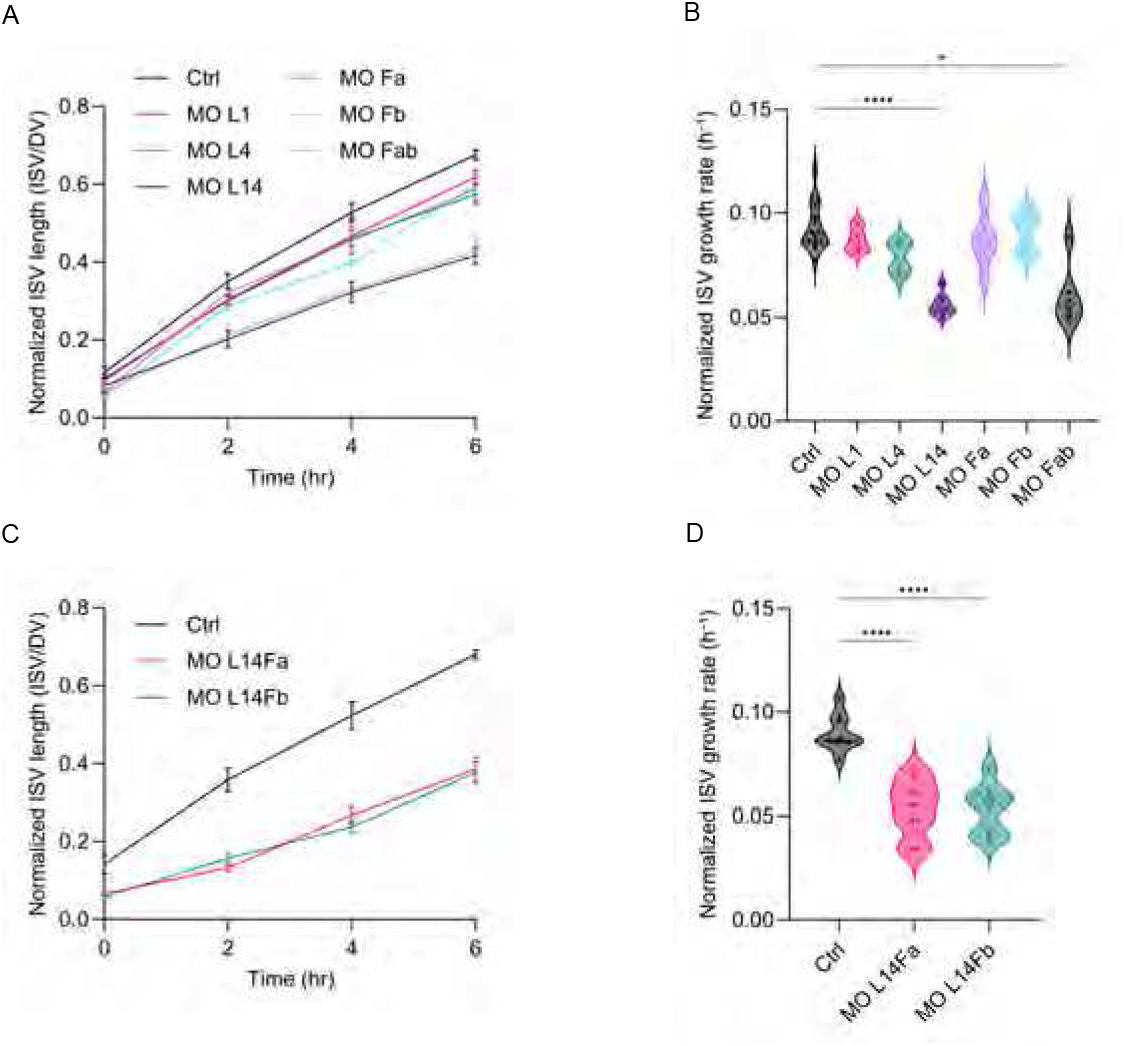
Dorsal–ventral axis normalization of ISV elongation and growth rates following ECM perturbation. (A) D/V-normalized ISV length over time in Control, MO L1, MO L4, MO L14, MO Fa, MO Fb, and MO Fab embryos. (B) D/V-normalized ISV growth rates for the same treatments. (C) D/V-normalized ISV length over time in Control, MO L14Fa, and MO L14Fb embryos. (D) D/V-normalized ISV growth rates for the combined laminin– fibronectin knockdown conditions. Time-lapse imaging was performed from 22 to 28 hpf, with time expressed relative to the start of imaging (0–6 h). Three ISVs were measured per embryo and averaged at each time point before normalization to the D/V-axis length of the corresponding embryo. Normalized growth rate was calculated as the change in normalized ISV length divided by the 6-h imaging interval (h^−1^). Length trajectories in (A,C) show mean ± s.e.m. Violin plots in (B,D) show distributions of embryo-level normalized growth rates, with individual dots representing embryos. For (A,B): Control *n* = 16, MO L1 *n* = 4, MO L4 *n* = 4, MO L14 *n* = 9, MO Fa *n* = 6, MO Fb *n* = 5, and MO Fab *n* = 6. For (C,D): Control *n* = 9, MO L14Fa *n* = 9, and MO L14Fb *n* = 8. Statistical comparisons of normalized growth rates were performed using a nonparametric Kruskal–Wallis test (B) and ordinary one-way ANOVA (D), with multiple comparisons against Control as indicated. Asterisks denote significance: \**p* < 0.05, \*\*\*\**p* < 0.0001. All *n* values represent independent embryos.

**Supplementary Figure 5:**
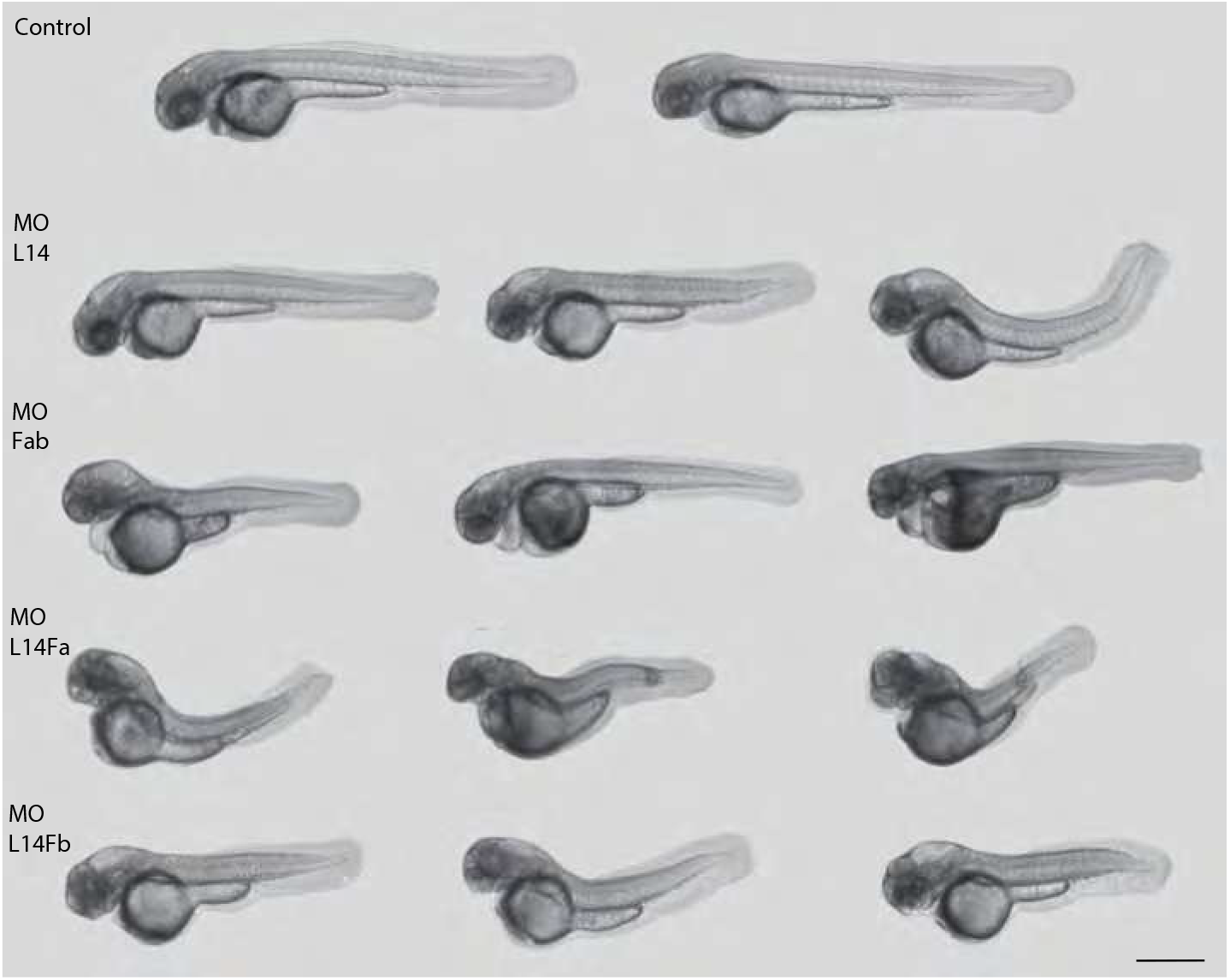
Whole-embryo morphology after laminin and fibronectin perturbation. Representative bright-field images of control morpholino-injected zebrafish embryos and embryos subjected to MO L14, MO Fab, MO L14Fa, or MO L14Fb perturbation at 48 hpf. These images were included to assess whether the vascular phenotypes tracked with gross developmental disruption across the principal combined ECM perturbation conditions. Gross trunk distortion is most evident in MO L14Fa, whereas MO L14Fb retains comparatively preserved overall trunk morphology despite clear vascular mispatterning. All embryos are oriented with anterior to the left and dorsal up. Scale bar, 500 *μ*m.

**Supplementary Figure 6:**
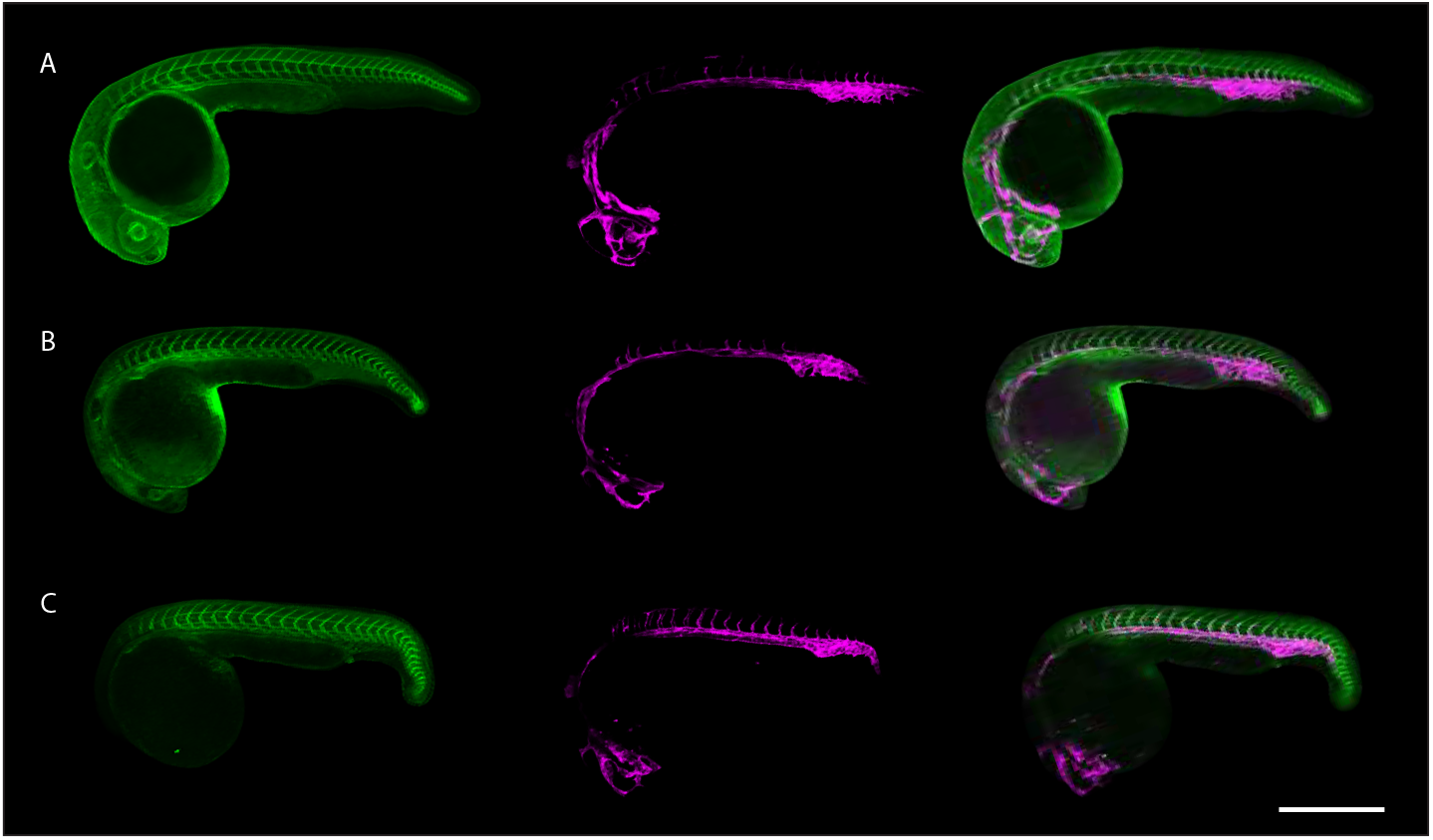
Spatial relationship between intersomitic ECM reporters and sprouting intersegmental vessels. Representative confocal images of double-transgenic zebrafish embryos during primary ISV outgrowth. Reporter combinations were imaged in separate experiments and are shown to illustrate spatial localization rather than quantitative differences in reporter abundance or developmental timing across lines. For each reporter combination, the ECM-reporter channel, endothelial channel, and merged view are shown. (A) *TgBAC(lama1:lama1-sfGFP)*;*Tg(kdrl:mCherry-CAAX)*, showing Lama1 reporter signal relative to sprouting endothelial cells. (B) *Tg(fn1a:fn1a-mNeonGreen)*;*Tg(kdrl:mCherry-CAAX)*, showing Fn1a reporter signal relative to sprouting endothelial cells. (C) *Tg(fn1b:fn1b-mCherry)*;*Tg(kdrl:EGFP)*, showing Fn1b reporter signal relative to sprouting endothelial cells. ECM-reporter signal is enriched in the intersomitic region adjacent to extending ISVs. Channels were pseudocolored consistently for visualization. All images are oriented with anterior to the left and dorsal up. Scale bar, 500 *μ*m.

**Supplementary Figure 7:**
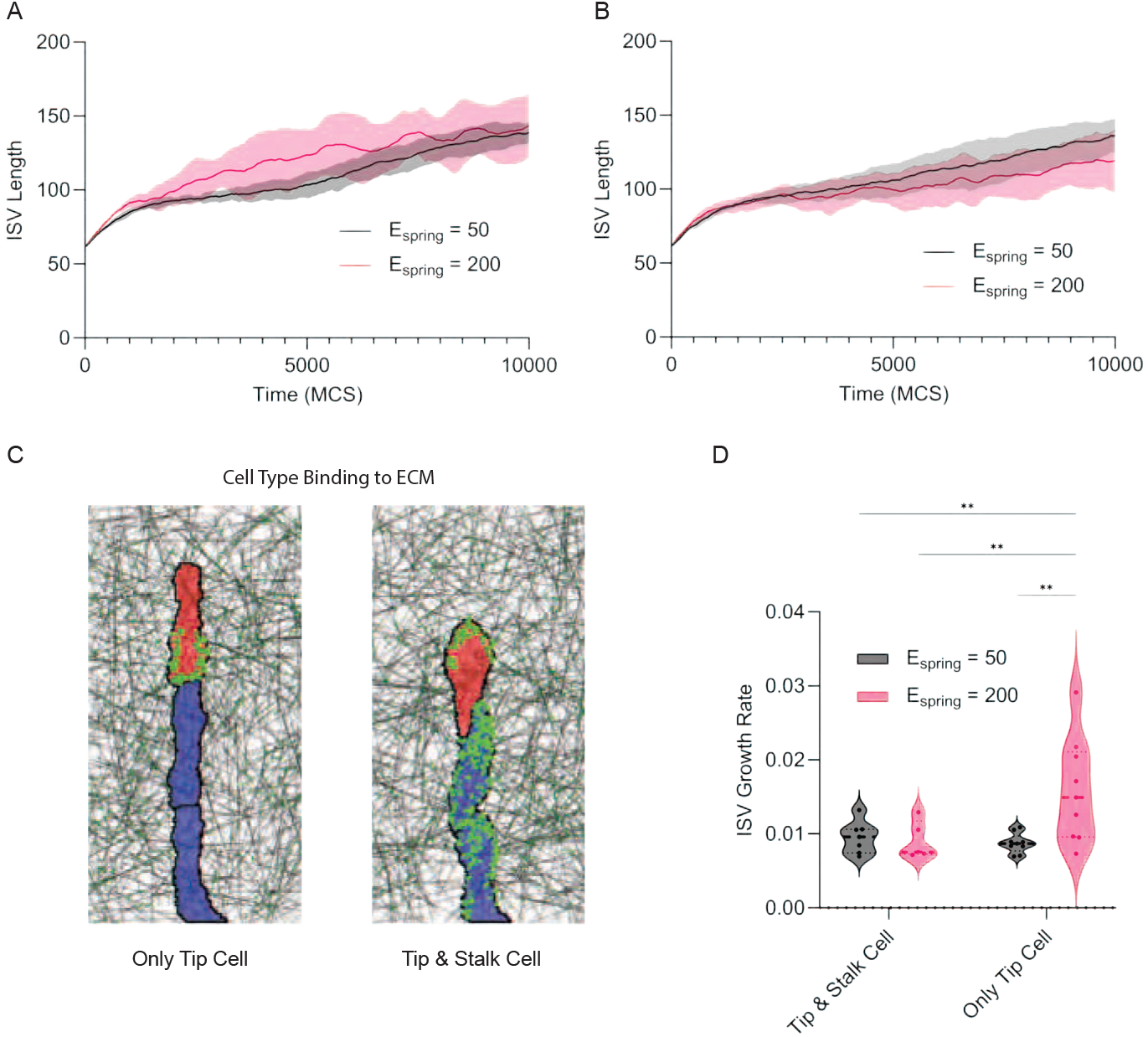
Effect of stalk-cell adhesion to the ECM. (A,B) Time series of vessel length when only the tip cell forms polarized FAs (A) or when stalk cells additionally form a spatially unpolarized distribution of FAs (B). (C) Simulation snapshot showing how FAs in stalk cells can immobilize the vessel and limit extension despite continued tip-cell migration. (D) Quantified vessel growth rate showing that stalk-cell FAs reduce vessel extension and eliminate sensitivity to ECM stiffness. This control tests whether stiffness sensitivity requires polarized tip-cell adhesion. The same simulation parameters were used as in Figure 3. Panels A, B, and D show mean and standard deviation over *n* = 10 simulations.

**Supplementary Figure 8:**
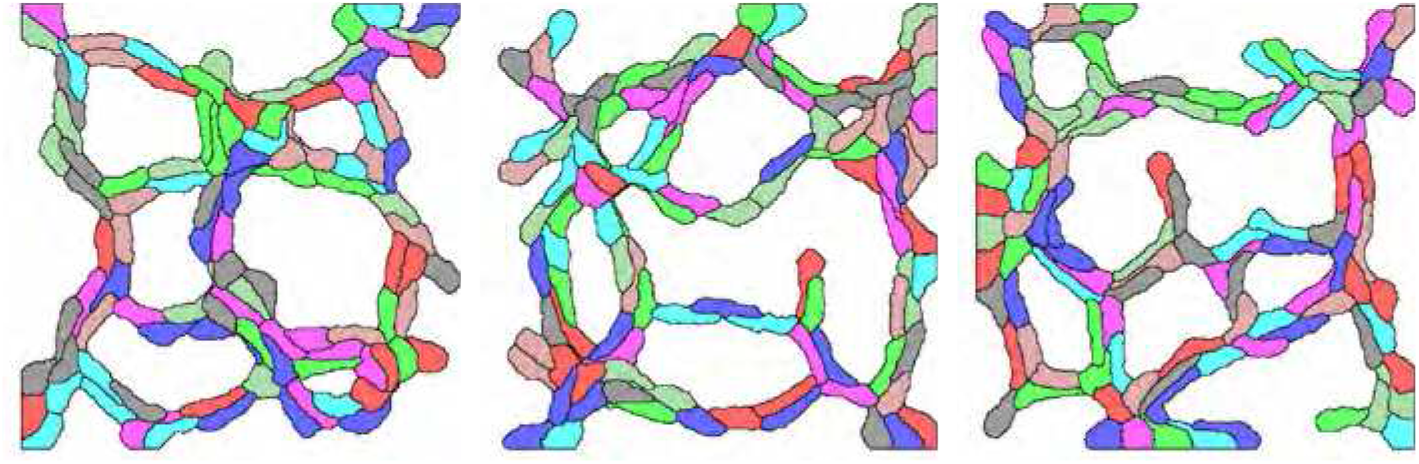
Simulations of the elongated cell model, a hybrid CPM model of endothelial network formation in which elongated endothelial cells are coattracted via a secreted chemoattractant (*12, 18*). Shown are three independent simulations of 100 cells in a 400 × 400 lattice.

**Supplementary Figure 9:**
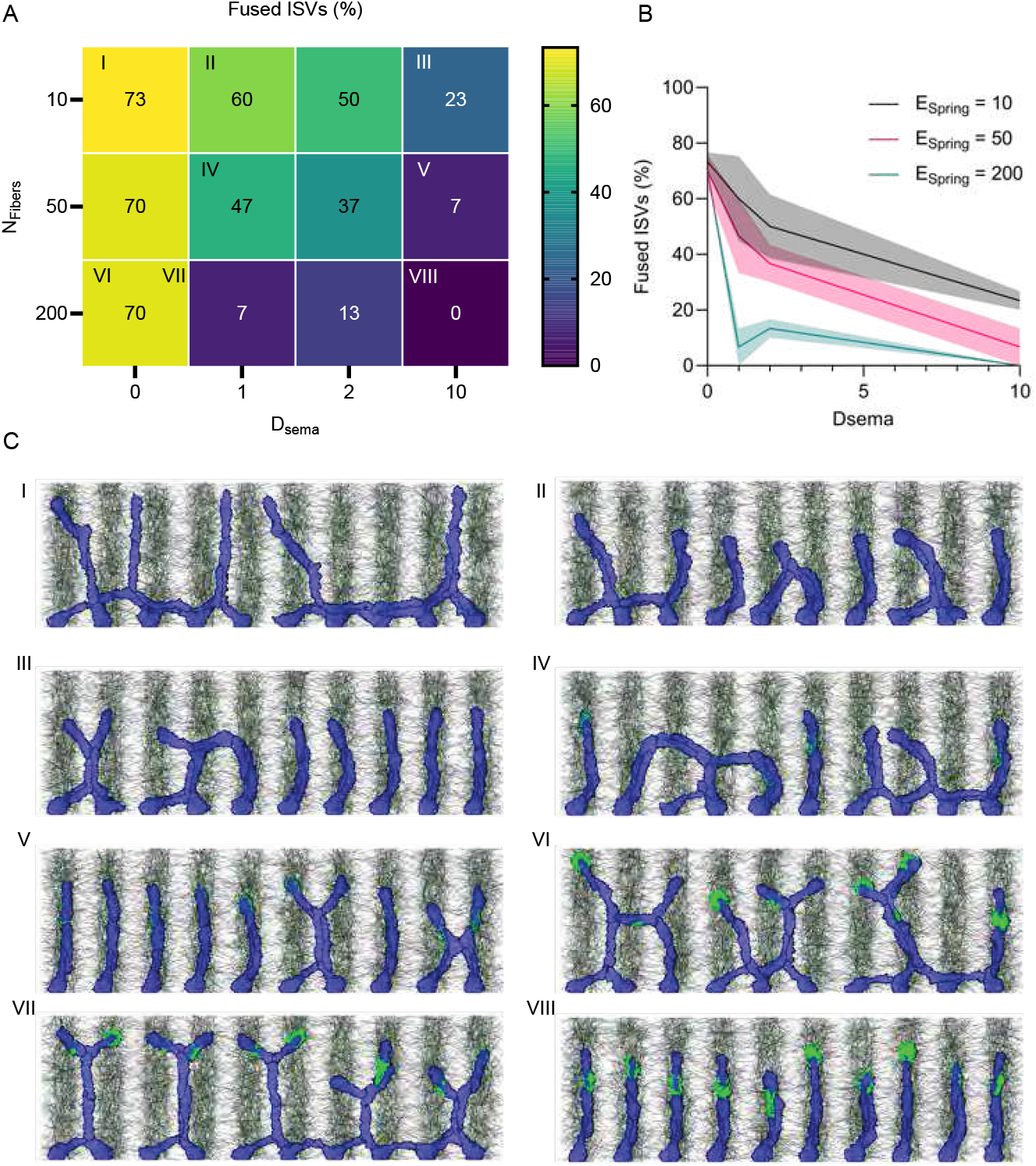
Simulations of ISV sprouting as a function of *E*_spring_ and *D*_sema_. A: Heatmap showing the percentage of fused vessels for combinations of *E*_spring_ and *D*_sema_ over *n* = 3 simulations for each condition. Low values indicate well-separated ISVs whereas high values indicate lateral fusion and network formation. B: Effect of *D*_sema_ on network formation. Average and standard deviation over *n* = 3 simulations for each condition. C: Representative simulation snapshots for the cases I-VIII indicated in panel A.

**Supplementary Figure 10:**
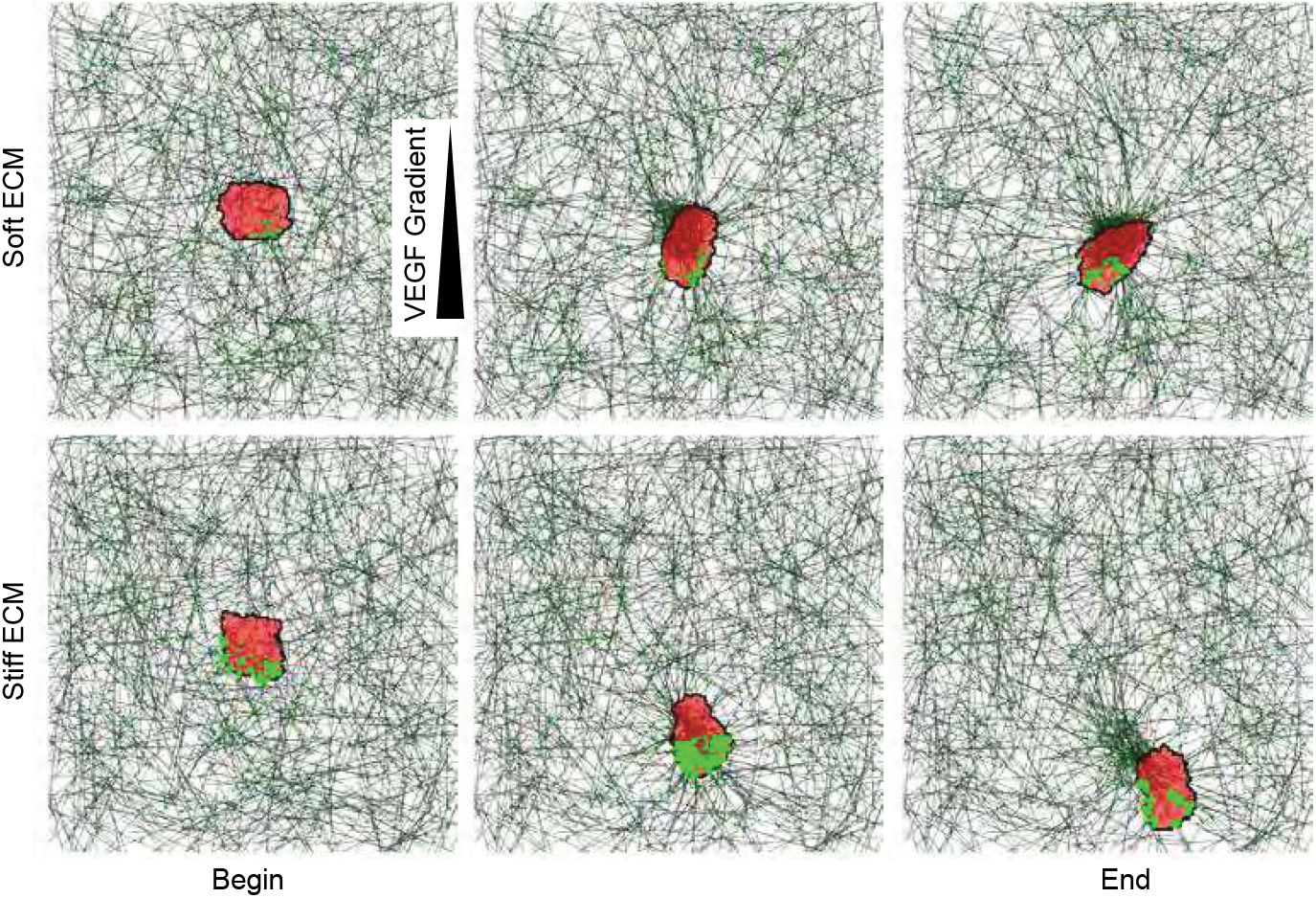
Snapshots of simulations of single-cell migration on ECM as a function of stiffness. The arrow indicates the VEGF gradient direction. Time progresses from left to right. On softer ECMs, cell migration is reduced, whereas on stiffer ECMs, the cell migrates directionally toward the VEGF source. The cell remodels both the softer and stiffer ECM, indicated by a degree of fiber densification at the rear of the cell and fiber reorientation around the cell. Fibers are drawn as gray lines and cross-links between fibers as green lines. The cell is shown in red and focal adhesions are shown as green discs. The size of the discs is proportional to the number of bound integrins in that FA.

**Supplementary Figure 11:**
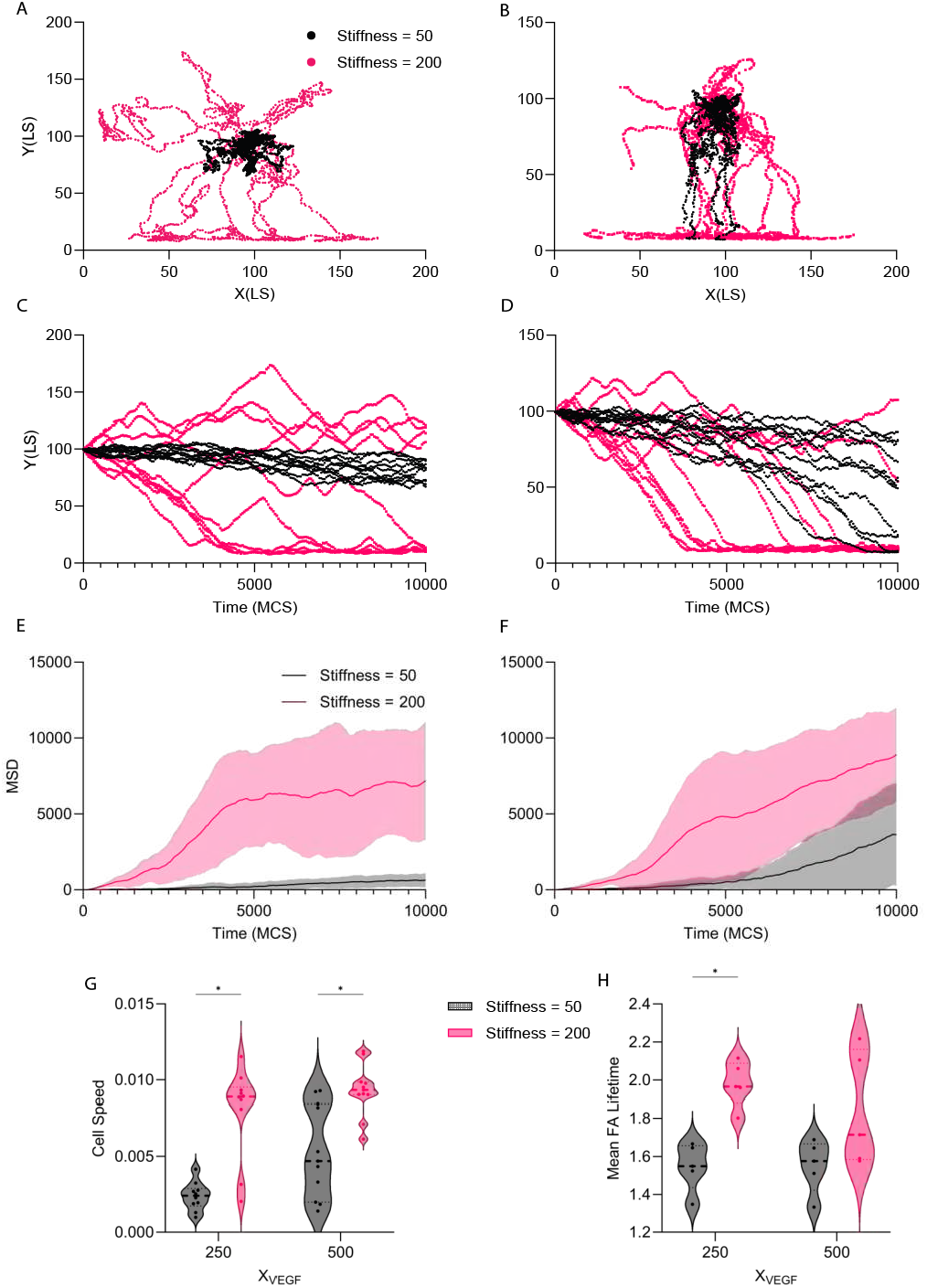
Analysis of single-cell migration simulations as a function of ECM stiffness and chemotactic sensitivity. (A,B) Trajectories of the cell center of mass for *χ*_VEGF_ = 250 (A) and *χ*_VEGF_ = 500 (B) under low (*E*_spring_ = 50) and high (*E*_spring_ = 200) ECM stiffness. (C,D) *y*-component of the cell center of mass as a function of simulation time for *χ*_VEGF_ = 250 (C) and *χ*_VEGF_ = 500 (D). (E,F) Mean squared displacement (MSD) as a function of simulation time for *χ*_VEGF_ = 250 (E) and *χ*_VEGF_ = 500 (F). Lines show the mean and shaded regions indicate ± s.d. across independent simulations. (G) Cell speed at 10,000 MCS as a function of chemotactic sensitivity and ECM stiffness. (H) Mean focal-adhesion (FA) lifetime per simulation as a function of chemotactic sensitivity and ECM stiffness. For (G,H), violin plots show the distributions and individual points represent independent simulations. Statistical comparisons between stiffness conditions at each *χ*_VEGF_ value were performed by ordinary two-way ANOVA followed by S^̌^idák’s multiple-comparisons test. Asterisks indicate the significant comparisons shown in the panels. Black indicates *E*_spring_ = 50 and magenta indicates *E*_spring_ = 200. For (A,C,E), *n* = 10 independent simulations per stiffness condition; for (B,D,F), *n* = 11 per stiffness condition. In (G), *n* = 10 per stiffness condition for *χ*_VEGF_ = 250 and *n* = 11 for *χ*_VEGF_ = 500; in (H), *n* = 5 independent simulations per condition.

## Supplementary Table

**Supplementary Table 1:**
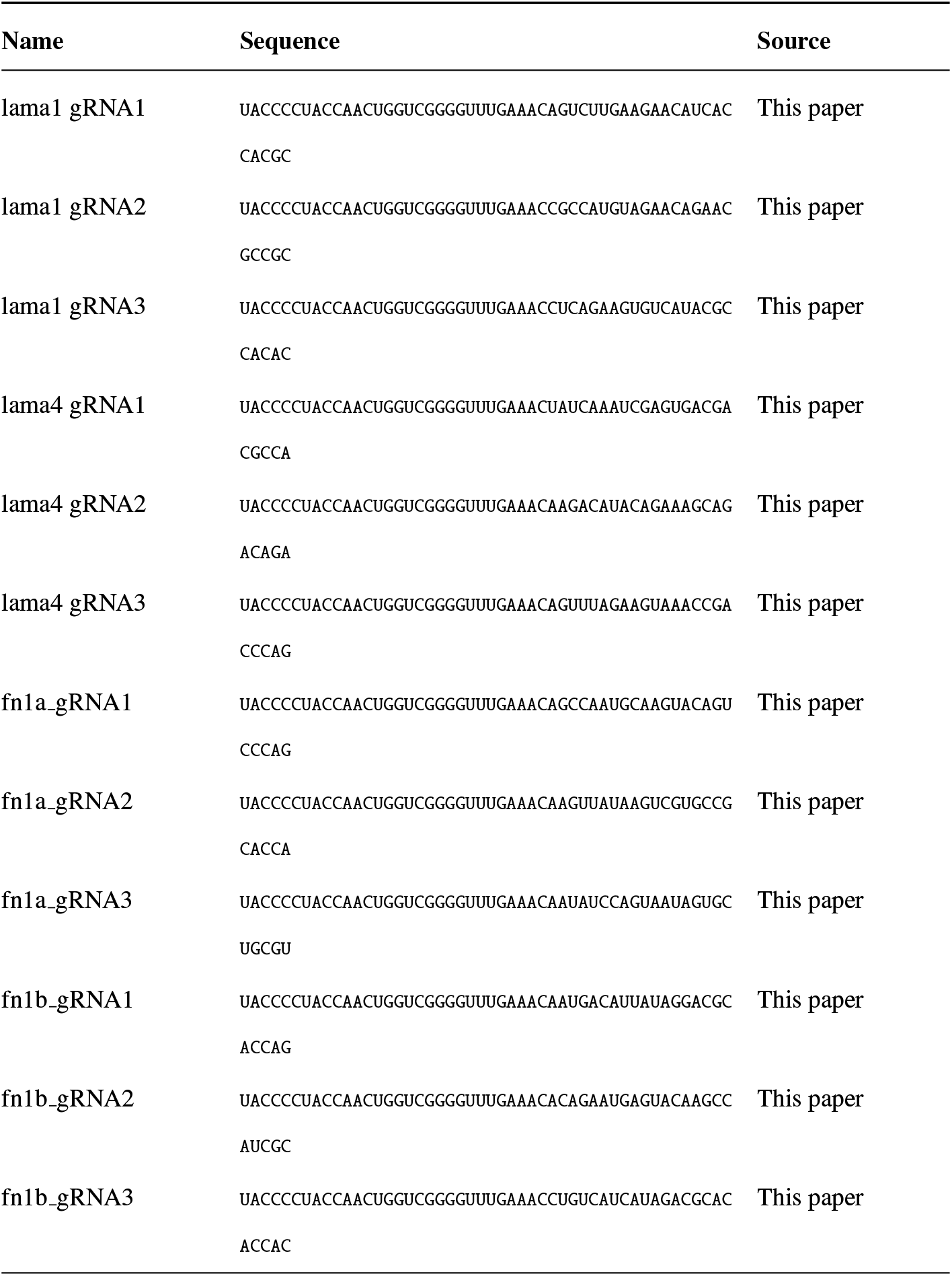
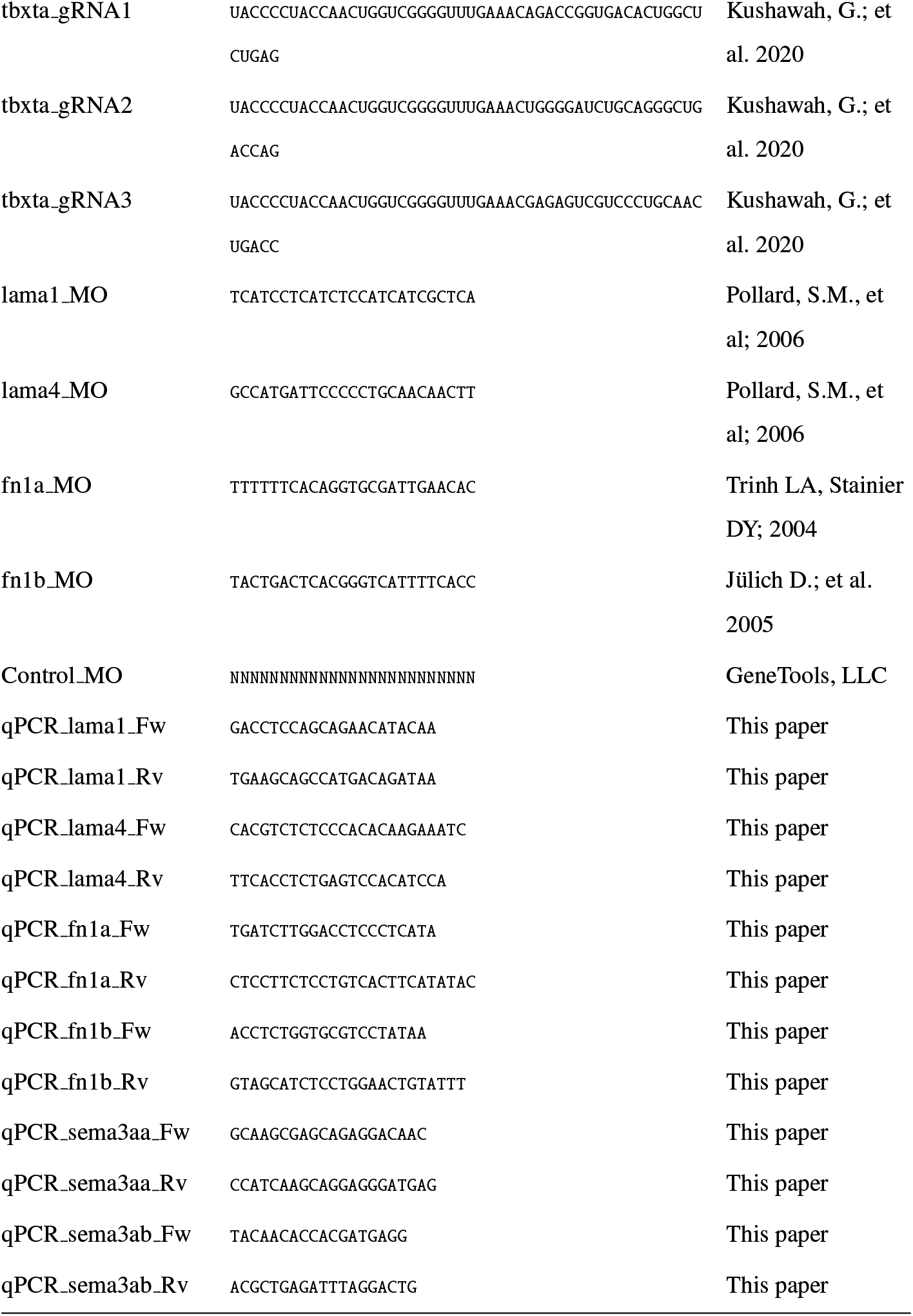
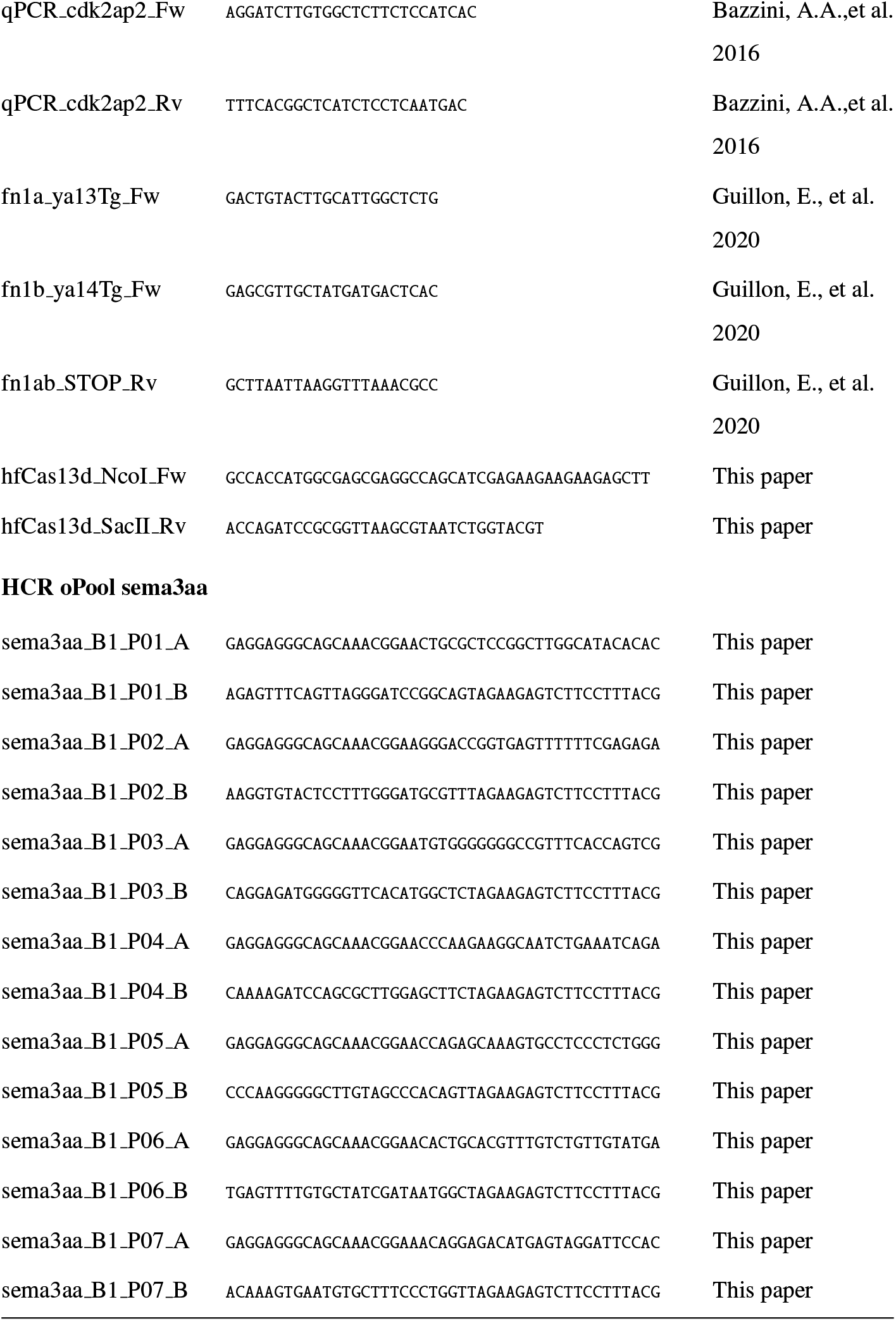
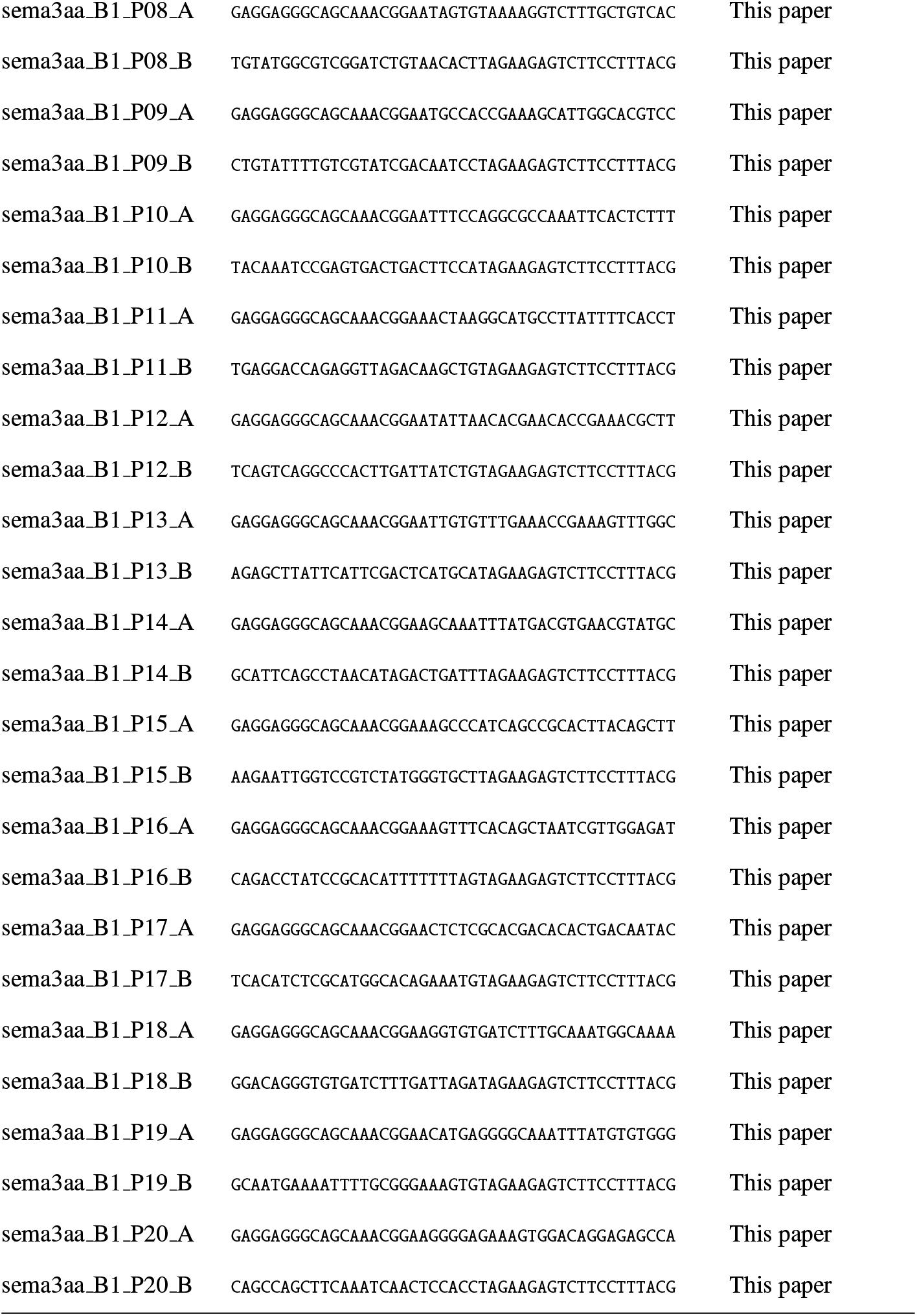
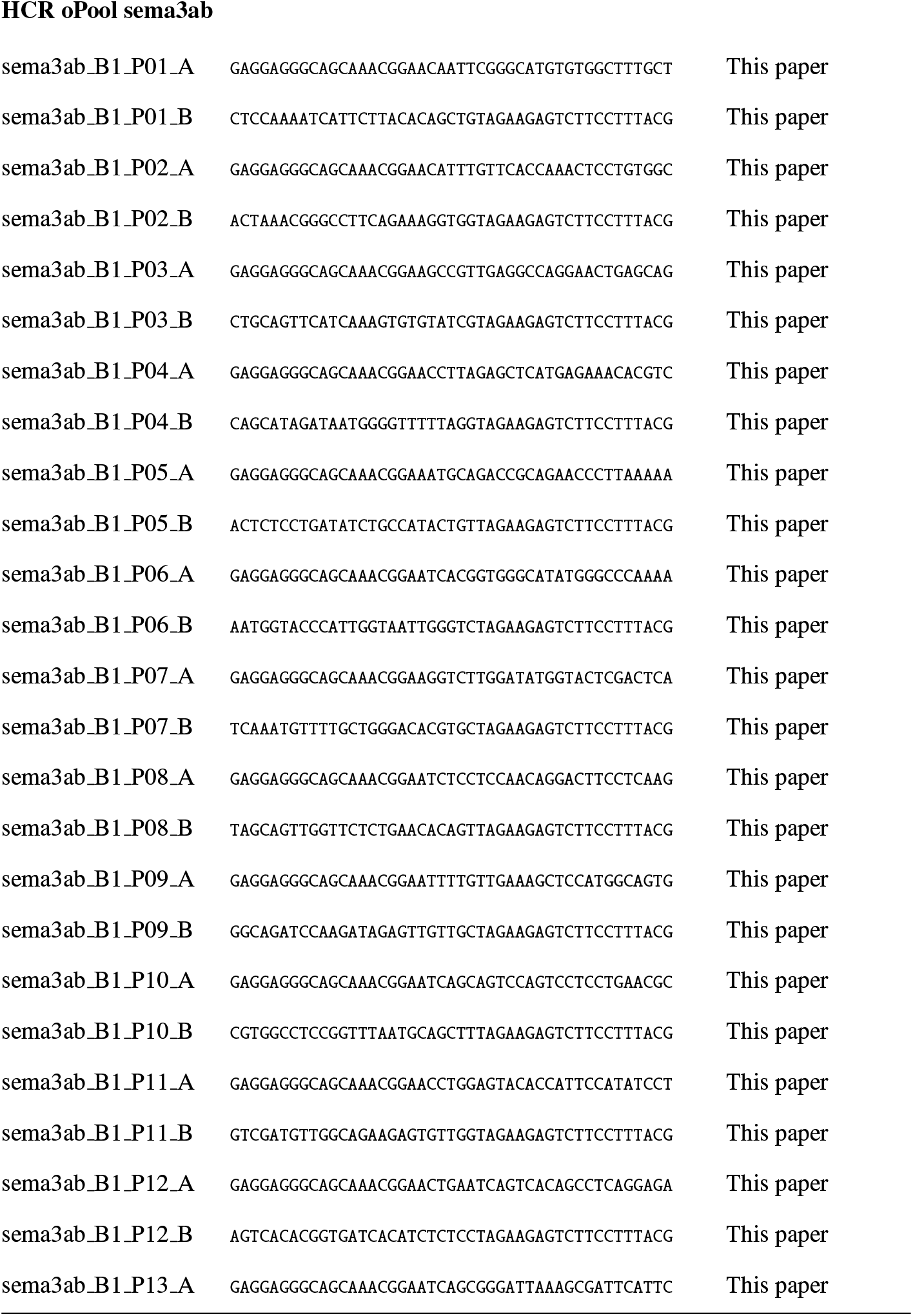
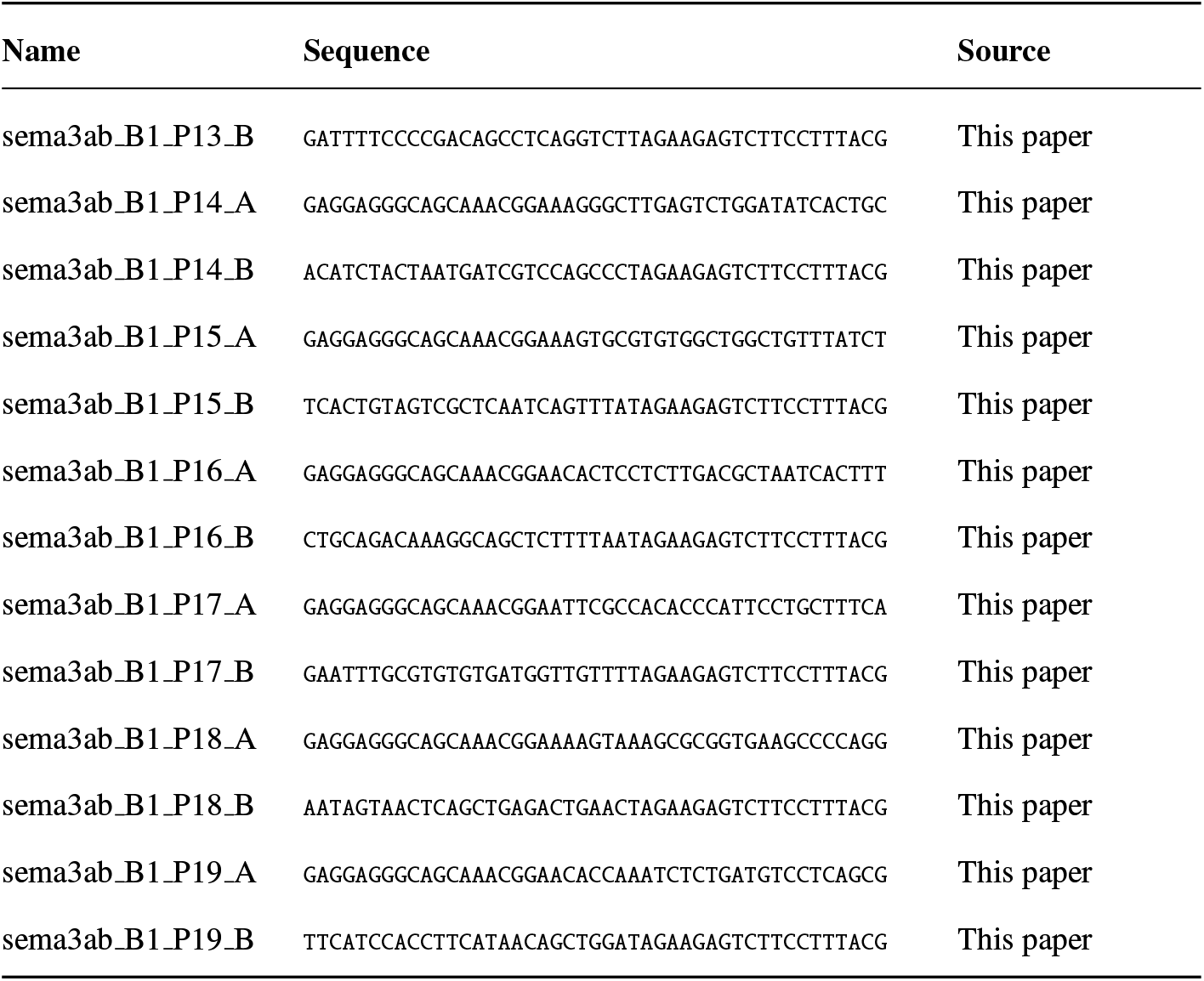
Sequences of morpholinos, qPCR primers, genotyping primers, gRNAs, and HCR oPools used in this study.

## Supplementary Videos

**Video 1:** 3D visualization of ISVs and laminin alpha 1. The movie opens with a brightfield 3D view of the zebrafish trunk at 24 hpf. Next, endothelial vessels are shown in magenta (*Tg(kdrl:mCherry-CAAX)*), followed by a green 3D visualization of laminin from the *lama1* reporter (*TgBAC(lama1:lama1-sfGFP)*). The entire 3D volume is created in Imaris from a confocal Z-stack to visualize enrichment within the intersomitic spaces. Orientation: anterior left, dorsal up. Scale bar, 10–15 *μ*m.

**Video 2:** 3D visualization of ISVs and fibronectin 1a. The movie opens with a brightfield 3D view of the zebrafish trunk at 24 hpf. Next, endothelial vessels are shown in magenta (*Tg(kdrl:mCherry-CAAX)*), followed by a green 3D visualization of Fn1a from the reporter (*Tg(fn1a:fn1a-mNeonGreen)*). The entire 3D volume is created in Imaris from a confocal Z-stack to visualize enrichment within the intersomitic spaces. Orientation: anterior left, dorsal up. Scale bar, 10–15 *μ*m.

**Video 3:** 3D visualization of ISVs and fibronectin 1b. The movie opens with a brightfield 3D view of the zebrafish trunk at 24 hpf. Next, endothelial vessels are shown in magenta (*Tg(kdrl:EGFP)*), followed by a green 3D visualization of Fn1b from the reporter (*Tg(fn1b:fn1b-mCherry)*). The entire 3D volume is created in Imaris from a confocal Z-stack to visualize enrichment within the intersomitic spaces. Orientation: anterior left, dorsal up. Scale bar, 10–15 *μ*m.

**Video 4:** Time-lapse visualization of an ectopic connection between neighboring intersegmental vessels following combined laminin and fibronectin perturbation (MO L14Fa) in *Tg(kdrl:EGFP)*. The embryo was imaged from 22 to 28 hpf. Endothelial fluorescence imaging shows two initially distinct ISVs extending along abnormal trajectories and subsequently establishing a direct lateral connection. This sequence provides direct evidence of an ISV fusion event and is therefore distinguished from other aberrant ISVs classified more broadly as ectopic branching, misguidance, or abnormal inter-ISV connections. Embryo orientation: anterior left, dorsal up. Scale bar, 50 *μ*m.

**Video 5:** Simulation of a single cell moving toward a chemokine source on soft ECM. Parameters are *χ*_chemokine_ = 100 and *E*_spring_ = 50.0.

**Video 6:** Simulation of a single cell moving toward a chemokine source on stiff ECM. Parameters are *χ*_chemokine_ = 100 and *E*_spring_ = 200.0.

**Video 7:** Simulation result of ISVs sprouting from the dorsal aorta over stiff ECM. A static Semaphorin guidance field between adjacent ISV trajectories helps to keep the sprouts directed. Parameters are *E*_spring_ = 200.0 and *N*_fibers_ = 600.

## Data Availability

**Code 1:** Simulation code and parameters used for all simulations in this manuscript are in https://github.com/mathbioleiden/Tissue-Simulation-Toolkit/tree/somite-ISV

**Dataset 1:** The image data belonging to this manuscript are available from https://10.6019/S-BIAD2920

